# A multicenter spatial transcriptomics atlas of human tuberculosis and non-tuberculous mycobacterial disease

**DOI:** 10.1101/2025.10.17.683002

**Authors:** Xun Jiang, Leonard Christian, Yuesi Xi, Liang Zhou, Ahmed Alaswad, Xiaoyi Zheng, Nienke van Unen, Lavinia Neubert, Jana Dietrich, Jan C. Kamp, Moritz Kayser, Mark Kühnel, Jens Hohlfeld, Hortense Slevogt, Marius M. Hoeper, Naftali Kaminski, Tobias Welte, Jan Fuge, Felix C. Ringshausen, Robert J. Homer, Danny D. Jonigk, Cheng-Jian Xu, Jonas C. Schupp, Yang Li

**Author notes:** Corresponding author: YL,; JCS,; CJX,. Tobias Welte, MD, died March 10th, 2024. Shared-first author (Xun Jiang, Leonard Christian). Shared-last author (Cheng-Jian Xu, Jonas C. Schupp, Yang Li).

## Abstract

Granulomas are the hallmark of mycobacterial (MB) infections, forming structured immune environments that contain bacteria but also drive disease persistence. However, their spatial and functional organization remains unclear. Using spatial RNA sequencing on 38 patient samples, we identified five distinct granuloma niches: a necrotic core, an immune-activated inner niche, an inflammatory and an extracellular matrix (ECM)-remodeling middle niche, an outer structural niche, and a tertiary lymphoid structure niche supporting antigen presentation. Immune activity peaks in the inner niche, transitioning to fibrosis at the periphery. Lymph node granulomas display reduced fibroblast involvement but stronger JAK-STAT activation. *Mycobacterium tuberculosis* (MTB) granulomas exhibit heightened JAK-STAT and IFN-γ signaling, while non-tuberculous mycobacteria (NTM) granulomas show increased hypoxia signatures. Compared to sarcoidosis, MB granulomas feature a structured adaptive immune response, marked by the clustering of plasma cells. Our findings, accessible via https://lab-li.ciim-hannover.de/mb-granuloma/, define key disease signatures, guiding biomarker discovery and therapeutic targeting in granuloma-related diseases.

## Introduction

Infections caused by mycobacteria (MB), including *Mycobacterium tuberculosis* (MTB) and non-tuberculous mycobacteria (NTM), are significant pulmonary health challenges with widespread global implications^1^. While MTB remains a leading cause of mortality in regions with high disease burden, NTM infections are increasingly recognized in both immunocompromised and immunocompetent populations^2^. Approximately two billion individuals globally are estimated to harbor latent mycobacterial infections, forming a reservoir for potential reactivation under conditions of immune compromise, co-infections or environmental factors^3–5^. Addressing these diseases requires a deeper understanding of MB-induced granulomas to improve prevention, diagnostics, and treatment strategies.

A hallmark of MB pathology is the formation of granulomas—organized aggregates of immune cells that attempt to contain the bacterial infection. In tuberculosis (TB), caused by MTB, infection begins when MTB-containing aerosols are inhaled, and alveolar macrophages engulf the bacteria^6^. MTB has evolved sophisticated mechanisms to evade host defenses, including inhibition of phagosome-lysosome fusion, allowing bacterial survival and replication within macrophages^7^. Additionally, MTB can suppress macrophage apoptosis by downregulating interleukin-10 (IL10) secretion and reducing tumor necrosis factor receptor 1 (TNF-R1) expression on the cell surface^8,9^. Over time, infected macrophages differentiate into so called “epithelioid cells”, which express high levels of adhesion molecules like E-cadherin, contributing to granuloma structure^10^. Following intracellular replication, MTB may induce necrosis of infected macrophages, enabling bacterial escape and further dissemination^11^. Unlike MTB, many NTM species do not rely on host-to-host transmission but rather cause opportunistic infections, particularly in individuals with pre-existing lung conditions, immunodeficiency or immunosuppression^12–15^. Due to the relatively lower incidence of NTM infections and the broad spectrum of bacterial strains involved, detailed research on NTM pathogenesis remains limited.

Granulomas induced by MB infections are characterized by distinct cellular architectures. The central necrotic core surrounded by epithelioid cells that maintain granuloma integrity and limit bacterial dissemination^16,17^. Other immune cells, including T cells, B cells, natural killer (NK) cells, and fibroblasts, are recruited to the granuloma periphery, contributing to the immune response and structural stability^10,18^. The spatial arrangement and interactions among these diverse cell types critically influence infection outcomes. In particular, inflammatory signals in granulomas can induce replication stress and trigger the DNA damage response, promoting the formation of polyploid and multinucleated macrophages^19^, which may further support granuloma structure and persistence.

We hypothesize that spatially distinct niches within granulomas regulate immune responses and tissue remodeling, balancing pathogen containment with mechanisms exploited by mycobacteria for chronic persistence. Despite extensive research on the general structure and cellular composition of MB granulomas, the specific functions, spatial organization, and interactions of immune cells within the granuloma microenvironment remain unclear. Unlike traditional RNA sequencing, spatial transcriptomics retains the tissue’s native architecture, enabling a detailed mapping of gene expression to anatomical structures, whose effectiveness to describe lung immunity has been demonstrated before^20–22^. Our analysis identified five distinct niches within granulomas, each characterized by unique cellular interactions and molecular activities. By integrating our findings with the Human Lung Cell Atlas (HLCA v2)^23–25^, we enhanced the resolution of immune and stromal cell populations and provided insights into granuloma-specific immune and remodeling processes. Spatial pathway analysis revealed distinct functional profiles, with localized immune activation and extracellular matrix dynamics varying across granuloma regions. Cell-cell interaction analysis highlights key mediators, including collagen and SPP1 signaling in extracellular matrix remodeling, and MIF and CXCL interactions in immune cell recruitment and activation across granuloma niches. A layered structure was predominantly observed in lung specimens, while MB infected lymph nodes displayed diffuse necrotic patches, heightened JAK-STAT, INF-γ and reduced TGF-β, hypoxia activity compared to pulmonary granuloma. Comparative analysis with non-infectious sarcoidosis granulomas revealed shared and unique features of granulomatous inflammation, highlighting disease-specific mechanisms. This spatial atlas advances our understanding of MB granulomas, revealing critical disease signatures and offering a foundation for identifying biomarkers and therapeutic targets to address granuloma-related diseases and MB persistence.

## Results

### Spatial transcriptional profiling of MB granulomas across 38 samples from three medical centers

We analyzed 38 formalin-fixed, paraffin-embedded (FFPE) lung tissue samples from mycobacteria-infected patients recruited from three independent institutes: Hannover Medical School (Germany), University Hospital RWTH Aachen (Germany), and Yale School of Medicine (USA). The cohort consisted of 51.3% male patients with a median age of 59 years ranged from 21 to 81 years old (Table S1). Using the Visium technology (10 x Genomics), we profiled the spatial gene expression within granulomas, enabling mapping of cellular and molecular interactions. The study was structured into a discovery cohort (27 lung and five lymph node samples) for hypothesis generation and a replication cohort (six lung samples) conducted independently by another laboratory to confirm the key findings (Figure 1A). This approach provided a robust framework to identify conserved spatial and functional patterns across granulomas while accounting for inter-individual variation.

**Figure 1.**
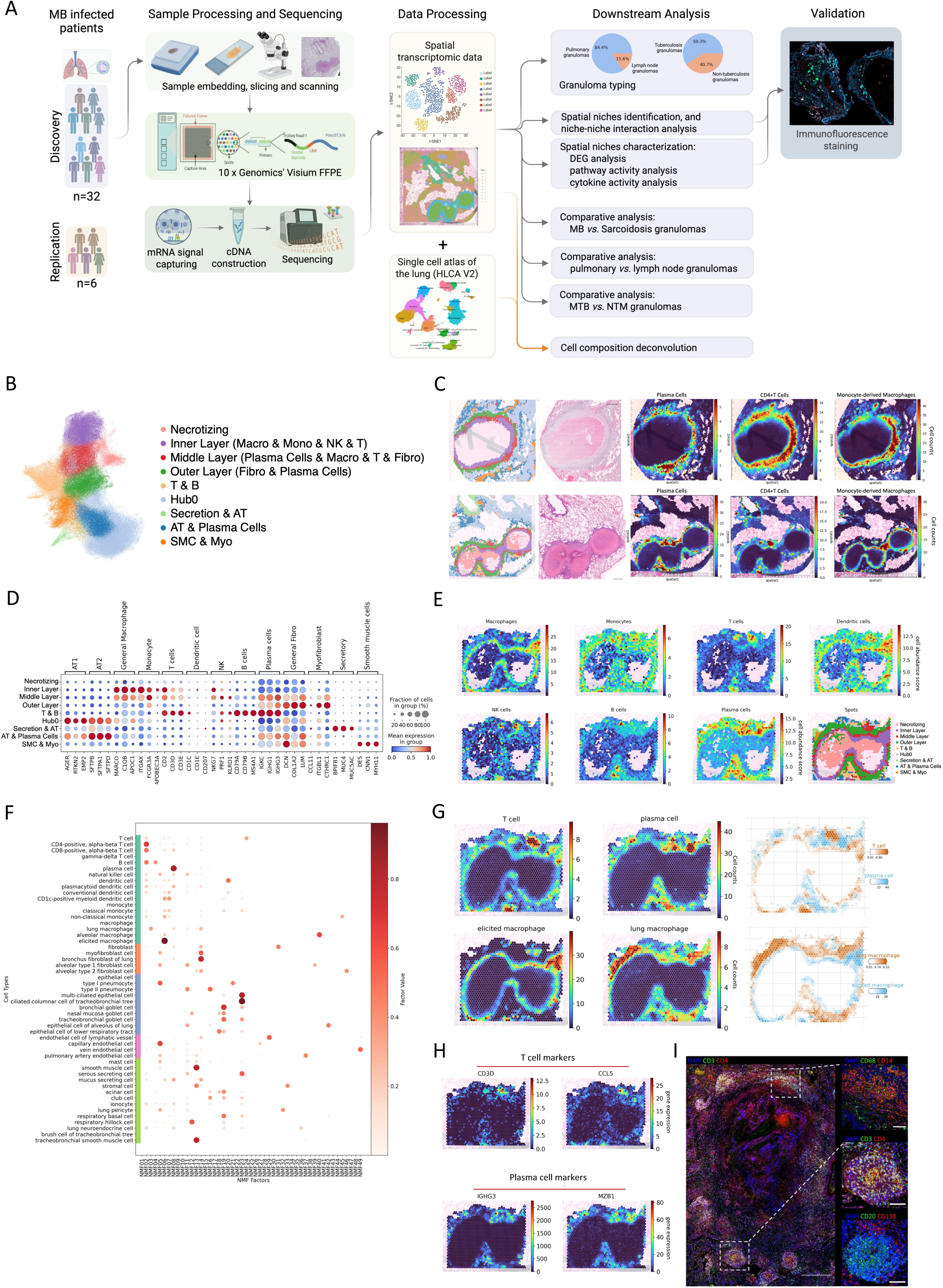
Spatial transcriptomic analysis of MB granulomas. (A) Schematic overview of the study workflow. Biobanked lung tissue samples were collected from a discovery cohort (n=32) and a validation cohort (n=6). The samples were processed using the Visium probe-based spatial transcriptomics (10x genomics). After H&E staining and cDNA library preparation, sequencing was performed to map spatial gene expression in granulomas. Spatial niches were identified using clustering and cell-type deconvolution methods, which were improved by integration of the Lung v2 dataset, leading to insights into the functional roles of each niche. All downstream analysis were listed in the graphic. Main findings from spatial transcriptomics were validated through immunofluorescence protein staining. (B) Spatial niche clustering and annotation based on integrated data from 32 granuloma samples. Distinct clusters representing necrotic, inner, middle, and outer niche, as well as other cellular regions, are visualized in UMAP. (C) Spatial annotation of niches is overlaid on histology images, validating correspondence with H&E staining and deconvoluted cell type distribution for each sample. (D) Correlation between annotated spatial niches and marker genes of key lung cell types. This association validates the accuracy of cell type deconvolution and spatial niche annotation. (E) Mapping of association scores between cell types and niches onto 2D spatial images to visualize cellular distributions within the granuloma structure. (F) Association between NMF factors and single-cell gene expression patterns. This figure shows the relationship between NMF factors derived from spatial transcriptomics samples and gene expression patterns from a single-cell reference dataset. Each row represents a specific cell type, categorized into major cell type clusters (immune cells, fibroblasts, epithelial cells, endothelial cells, and others), as indicated by the color-coded legend on the left. The size and color intensity of the dots indicate the strength of the association (factor value) between a given NMF factor (x-axis) and the gene expression profile of a specific cell type (y-axis). (G) Differential distribution of T cells, plasma cells, and macrophages in the granuloma identified using Cell2location. This figure shows the spatial distribution of T cells, plasma cells, monocyte-derived macrophages, and interstitial macrophages within the granuloma region, inferred using the Cell2location package. T cells and plasma cells are predominantly localized in lymphocytic regions of the granuloma, while monocyte-derived macrophages (elicited) are concentrated near the granuloma core, alveolar macrophages (lung) are located in the middle niche, highlighting the compartmentalized nature of immune and structural cell niches. (H) Visualization of T cell and plasma cell marker gene expression in MB granuloma. The left panels show the expression of T cell markers CD3D and CCL5 (left to right), indicating regions enriched in T cells, predominantly localized to lymphocytic areas. The right panels display plasma cell markers IGHG3 and MZB1 (left to right), highlighting plasma cell-enriched zones that align with the granuloma’s peripheral lymphocytic regions. (I) Representative immunofluorescence protein staining of a MB granuloma shows the presence of monocyte-derived macrophages in the inner niche (CD68, green; CD14, red; upper panel), T cells in the middle and T&B niche (CD3, green; CD4 red; large and middle panel), as well as plasma and B cells in the T&B niche (CD20, green; CD138, red; lower panel); Nuclei stained with DAPI (blue); Scale bar: 500 µm (large panel) and 100 µm (small panel).

A total of 301,656 spots were detected under the tissue, with a median of 6,151 unique molecular identifiers (UMI) per spot across the samples (Figure S1A), and with a median of 3,708 unique genes per spot (Figure S1B). Detailed information of sample quality was provided in Table S2. We observed consistent average gene counts and transcript levels across samples, indicating high data quality. The raw sequencing data has been deposited at European Genome-phenome Archive (waiting approval), and the processed data can be explored at the supplemental website https://lab-li.ciim-hannover.de/mb-granuloma/.

### Gene expression patterns define five distinct granuloma niches with specialized immune and structural functions

To investigate the spatial organization of MB granulomas, we integrated Leiden clustering of spatial gene expression with histological features, identifying five distinct granuloma niches (Figure 1B). The innermost cluster, the necrotizing niche, is characterized by low overall gene expression and a lack of specific cell type markers, indicative of extensive cell death and tissue destruction. Immediately surrounding the necrotic core niche is the inner niche, which is enriched in monocyte-derived macrophages, monocytes, and T cells, with dendritic cells and NK cells scattered in, reflecting active immune engagement with the pathogen. The next layer of the granuloma, the middle niche, contained a mixed population of T cells, plasma cells, macrophages, and fibroblasts. The outermost niche is dominated by fibroblasts, with some plasma cells mixed in. In addition, we identified distinct cell clusters that were often integrated into the outer niche comprising T, B and plasma cells (T&B niche), resembling tertiary lymphoid structures (Figure 1C, 1D and 1E). Collectively, these spatially defined niches reveal a layered organization within MB granulomas, reflecting a specialized compartmentalization in pathogen containment, immune activation, and tissue remodeling.

To validate our spatial annotation and refine cell composition estimates within MB granulomas, we conducted a cell composition deconvolution and identified non-negative matrix factors (NMF) based on covarying gene expression^26^ (Figure 1F). The results showed that the cell distributions aligned closely with the previously defined granuloma niches. T cells were predominantly localized in the inner granuloma niches, reinforcing their role in active immune engagement (Figure 1G, and S1B, NMF 2 and NMF 24). In contrast, plasma cells were enriched in the outer and middle niches, suggesting their involvement in antibody-mediated responses (Figure 1G, and S1B, NMF 8). Such cell distributions were further confirmed by the T and plasma cell marker genes’ expression (Figure 1H). Notably, macrophages dominated the inner niche, but their subtypes exhibited distinct spatial distributions. Monocyte-derived macrophages were localized near necrotizing and immune-active regions (Figure 1G, and S1B, NMF 6). In contrast, the alveolar macrophages were more evenly distributed in peripheral tissues (Figure 1G, and S1B, NMF 6 and NMF 40). These findings highlight the dynamic recruitment of monocyte-derived macrophages in response to infection. To further validate these results on a protein level (Figure 1I), we performed immunofluorescence protein staining of marker genes for the main cell types of the granuloma, which matched the results of the deconvolution.

To further characterize niche-specific molecular functions, we performed differential gene expression (DEG) analysis to identify marker genes unique to each niche (Figure 2A, and S2A). The inner niche was abundant in *SPP1+* macrophages, prominent in idiopathic pulmonary fibrosis (IPF)^27^. The expression of *MMP9* and *MMP12* in this niche suggests a role of these macrophages in extracellular remodeling. In the inner and middle niche, lysosomal genes such as *LYZ*, *CTSB* and *CTSZ* were highly expressed, reflecting active pathogen clearance. The expression of the fibroblast-associated genes *COL1A1, COL3A1,* and *SPARC* in the outer niche highlights ECM remodeling around the granuloma. This niche is particularly enriched with *CTHRC1*+ fibroblasts, which have been linked to pulmonary fibrosis as well^28,29^. The T&B niche demonstrated elevated expression of lymphocyte markers, including *MS4A1*, *BLK*, *CD79A* and *CD79B*, along with *FCMR*, *CD37*, and *SELL*, indicative of lymphocyte differentiation, migration and adhesion (Figure 2A). Our findings demonstrate that MB granulomas exhibit a highly structured immune architecture, with spatially distinct niches specializing in immune defense and structural remodeling.

**Figure 2.**
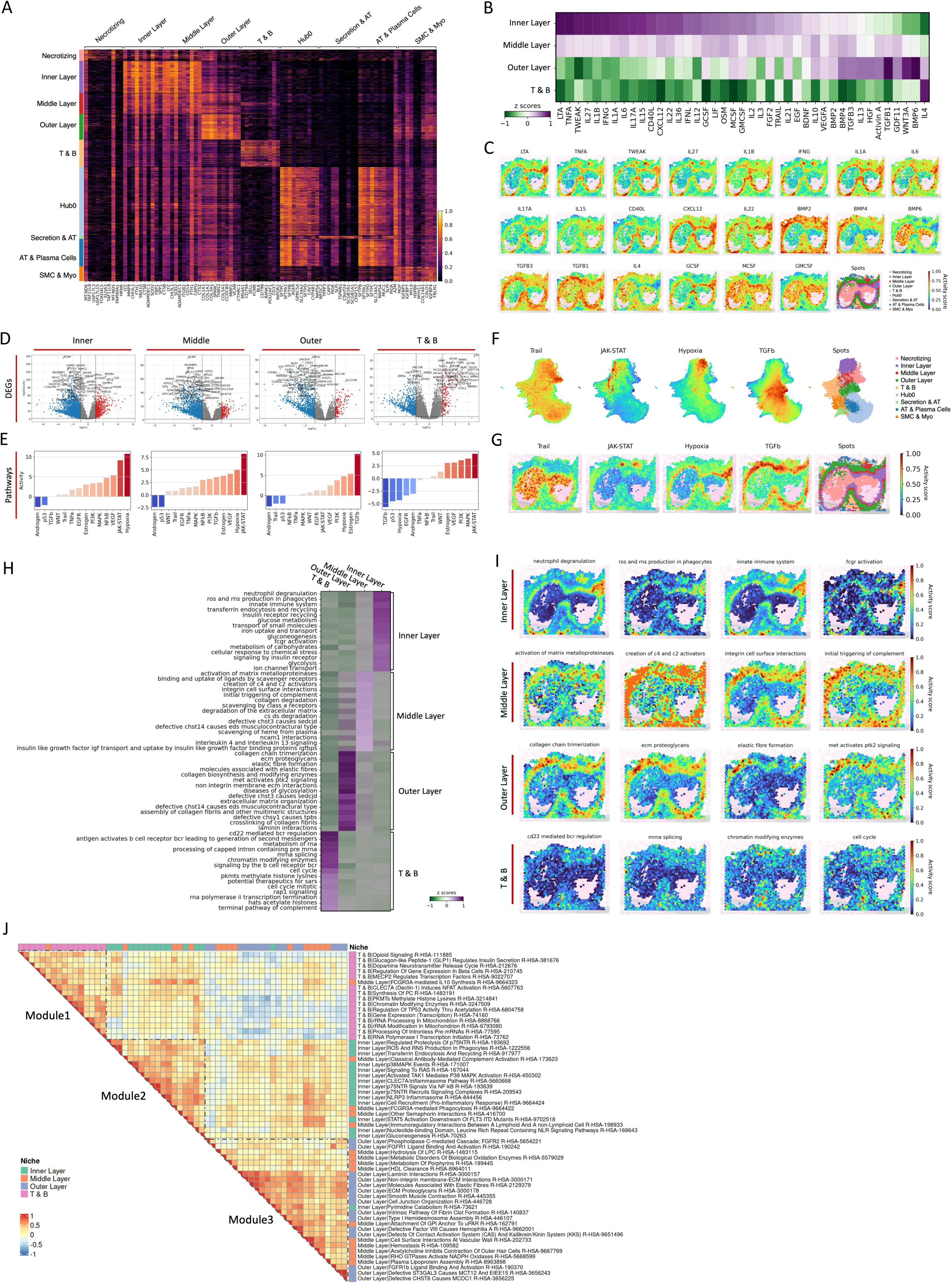
Cytokine activity and pathway dynamics in MB granulomas. (A) Top 10 DEGs from each Niche. (B) Cytokine activity Scores Across Granuloma Niches Inferred by MLM Analysis based on the CellTypist database. This heatmap illustrates Z-scores for distinct granuloma niches. Warmer colors (purple) indicate higher pathway enrichment scores, while cooler colors (green) represent lower activity. (C) Spatial Mapping of Cytokine Activity. (D) Pseudo-Bulk RNA-seq Analysis Revealing DEGs and (E) Pathway Activity Profiles for Granuloma Niches. The volcano plots show DEGs comparing each niche to the rest, with upregulated genes in red and downregulated genes in blue. (F) Mapping of Pathway Activity Using UMAP. This figure visualizes the distribution of key signaling pathways, including TRAIL, JAK-STAT, TGF-β, and Hypoxia, across MTB granulomas using UMAP embedding. (G) Spatial Mapping of Pathway Activity. (H) Enrichment scores of REACTOM pathways across granuloma niches. (I) Spatial Mapping of REACTOM Pathway Activity as an example. (J) Pathway Correlation and Modular Analysis Across Granuloma Niches. This heatmap shows pathway correlations across granuloma niches (inner niche, middle niche, outer niche, T&B niche). Three modules were identified: Module 1, enriched mainly in T & B cell activities; Module 2, featuring pro-inflammatory and innate immune responses mainly in the inner niche; and Module 3, associated with structural and metabolic processes in the outer niche.

As a variety of cytokines play a vital part in granuloma formation, we next examined the cytokine activity level across the granuloma to assess niche-specific involvement in cell recruitment, immune activity and tissue remodeling, based on the gene expression profile^30,31^ (Figure 2B). The inner niches of the granuloma exhibit hyperactivity of LTA, TNF-α, TWEAK, IL27, IL1B, IFN-γ, IL1A, IL6, IL17A, IL15, CD40L, CXCL12, and IL22, reflecting a highly pro-inflammatory and immune-dominant microenvironment. This cytokine profile indicates intense macrophage and T cell activation (TNF-α, IFN-γ, IL-1 family), robust neutrophil recruitment (IL-6, IL-17A), and enhanced tissue remodeling signals (TWEAK, CXCL12, CD40L). The high specificity of GCSF (CSF3) in the inner niches suggests a strong role in neutrophil recruitment and activation^32^, reinforcing the innate immune response within the granuloma core. While these factors support macrophage and neutrophil-driven pathogen defense in the inner niche, they also facilitate macrophage polarization and extracellular matrix remodeling in the outer niche, contributing to both immune containment and fibrosis. The high activity of MCSF (CSF1) and GMCSF (CSF2) in the inner niche and necrotizing region reflects the ongoing activation of monocytes and macrophage-driven^33^. In contrast, the middle niche showed reduced cytokine activity. The outer niche demonstrated a contrasting cytokine profile characterized by elevated levels of BMP2, BMP4, BMP6, TGF-β3, TGF-β1, and WNT3A, which are essential for extracellular matrix remodeling and tissue repair. These processes reflect the outer niche’s structural and reparative functions, encapsulating the granuloma while maintaining its integrity. Notably, the T&B niche exhibited high activity of IL4, indicative of a Th2 response contributing to immunomodulatory activity^34^, supporting adaptive immune processes such as antibody production and T cell regulation (Figure 2B, and 2C). This comprehensive study of these spatially organized immune dynamics will provide a framework for targeted therapeutic interventions that enhance pathogen clearance while preserving tissue function.

### Pathway activity gradients reveal functional compartmentalization within MB granuloma niches

To investigate the signaling pathway activities across MB granuloma niches, we performed pathway activity analysis using pseudo-bulk data. The inner niche displayed significant activation of JAK-STAT and hypoxia signaling pathways (Figure 2D, and 2E), indicating a hypoxic environment and robust immune activity aimed at pathogen containment^35–38^. The middle niche, containing plasma cells and macrophages, exhibited similar pathway activity patterns to the inner niche, albeit at reduced levels (Figure 2E). This suggests that the middle niche functions as a transitional zone, sustaining immune engagement with less intensity. In contrast, the outer niche, predominantly composed of fibroblasts, exhibited elevated TGF-β signaling (Figure 2E), reflecting its role in ECM remodeling. The T&B niche showed high JAK-STAT, but low TGF-β activity, probably due to its dominant immune cells (Figure 2E).

To further examine the spatial dynamics of pathway activities in MB granulomas, we compared adjacent niches — inner *vs.* middle and middle *vs.* outer niches (Figure S2B). Hypoxia-responsive and JAK-STAT signaling peaked in the inner niches, reflecting active immune responses and pathogen containment, while these activities decreased in the middle and outer niches. Conversely, TGF-β activity increased toward the outer niches, emphasizing a functional shift to ECM remodeling. We were able to validate these findings by applying multivariate linear model (MLM) analysis to infer pathway enrichment scores for each spatial spot, which corroborated the pseudo-bulk analysis results (Figure 2F, and 2G). Expression patterns of marker genes colocalized with these pathways, providing additional evidence for the functional roles of specific niches (Figure S2C).

To elucidate the functional differences across the granuloma niches, we performed a pathway analysis using the Reactome (Figure 2H, and 2I) and KEGG databases (Figure S2D). The inner niche exhibited robust immune activity characterized by neutrophil degranulation, production of reactive oxygen and nitrogen species (ROS and RNS) in phagocytes, innate immune system activation, and Fc gamma receptor-mediated signaling. The inner niche also displayed a shift in energy metabolism towards carbohydrates supporting the immune reaction as presented by high glyconeogenesis, glycolysis and glucose metabolism pathway activity. These processes highlight the inner niche’s role in mounting a strong innate immune response to contain the pathogen under hypoxic conditions. Across the granuloma niches, the middle niche showed the highest activity of scavenger receptor-related pathways, which have been implicated to support granuloma chronicity^39^. Serving as a transitional zone, the middle niche showed both activation of the complement system (C4 and C2 activators) and pathways involved in matrix metalloproteinase activity, and integrin-mediated cell surface interactions. In contrast, the outer niche displayed enrichment in processes related to ECM remodeling, including collagen chain trimerization, ECM proteoglycan deposition, elastic fiber formation, and signaling pathways such as MET activation and PTK2 signaling. The granuloma-associated T&B cell niche demonstrated adaptive immune processes, including CD22-mediated B cell receptor regulation and signaling by B cell receptors, underscoring their contributions to antigen-specific immune responses. This spatial distribution of biological processes underscores the granuloma’s functional complexity, with the inner niche focusing on immune defense, the middle niche serving as a mediator between immune and structural roles, and the outer niche specializing in structural maintenance.

### Niche-specific pathway dynamics reveal a balance between immune activation and tissue remodeling in MB granuloma

To better characterize the spatially functional compartmentalization of the granuloma niches, we examined their correlation using AUC scores between pathways in the inner, middle, outer, and T&B niches (Figure 2J). Unsupervised clustering revealed three distinct modules, each corresponding to specific biological processes within the granuloma’s spatial compartments. Module 1, enriched in the T&B niche, reflects transcriptional and epigenetic processes, such as PKMT-mediated histone methylation, TP53 regulation, and mitochondrial rRNA modification, that may support cytokine production and antigen presentation in adaptive immune regulation. Module 3, associated with the outer granuloma niche, highlights structural and metabolic adaptations, including extracellular matrix remodeling (e.g., laminin interactions, FGFR signaling) and lipid metabolism (e.g., HDL clearance, lipoprotein assembly), reflecting tissue repair mechanisms^40–42^. Interestingly, Module 1 is negatively correlated with Module 3, suggesting an inverse relationship between adaptive immune activities and structural/metabolic processes, likely reflecting their spatial and functional segregation. Module 2, representing pro-inflammatory innate immune responses in the inner granuloma niche, exhibits weak positive correlations with both Modules 1 and 3. Additionally, pathways within the same niches generally showed positive associations, except for the middle niche, where pathways were linked with other niches, emphasizing its unique mediatory role in coordinating granuloma functionality. These findings underscore the granuloma’s spatial compartmentalization and the dynamic interplay between immune and metabolic processes, suggesting a finely tuned balance between immune activation and tissue remodeling.

### Immune and structural interactions shape granuloma architecture

To elucidate how the granuloma niches communicate with each other, we conducted a niche-niche interaction analysis. The interaction strength between different granuloma niches demonstrated robust communication between the inner, middle and outer niches (Figure 3A), with the necrotizing niche exhibiting limited outgoing signals and an absence of incoming interactions, suggesting a localized microenvironment largely isolated from external regulation. In contrast, the inner and middle niches demonstrate both strong outgoing and incoming signaling activity of immune molecules, such as ICAM, SPP1 and MIF, indicating their central role in orchestrating immune responses and extracellular matrix remodeling. The outer niche shows moderate outgoing signaling but relatively lower incoming activity, indicating its involvement in structural maintenance and remodeling rather than direct immune regulation, while the T&B niche primarily functions as a recipient of signals, integrating immune cues from other niches to sustain adaptive immune responses within the granuloma (Figure 3B, and 3C).

**Figure 3.**
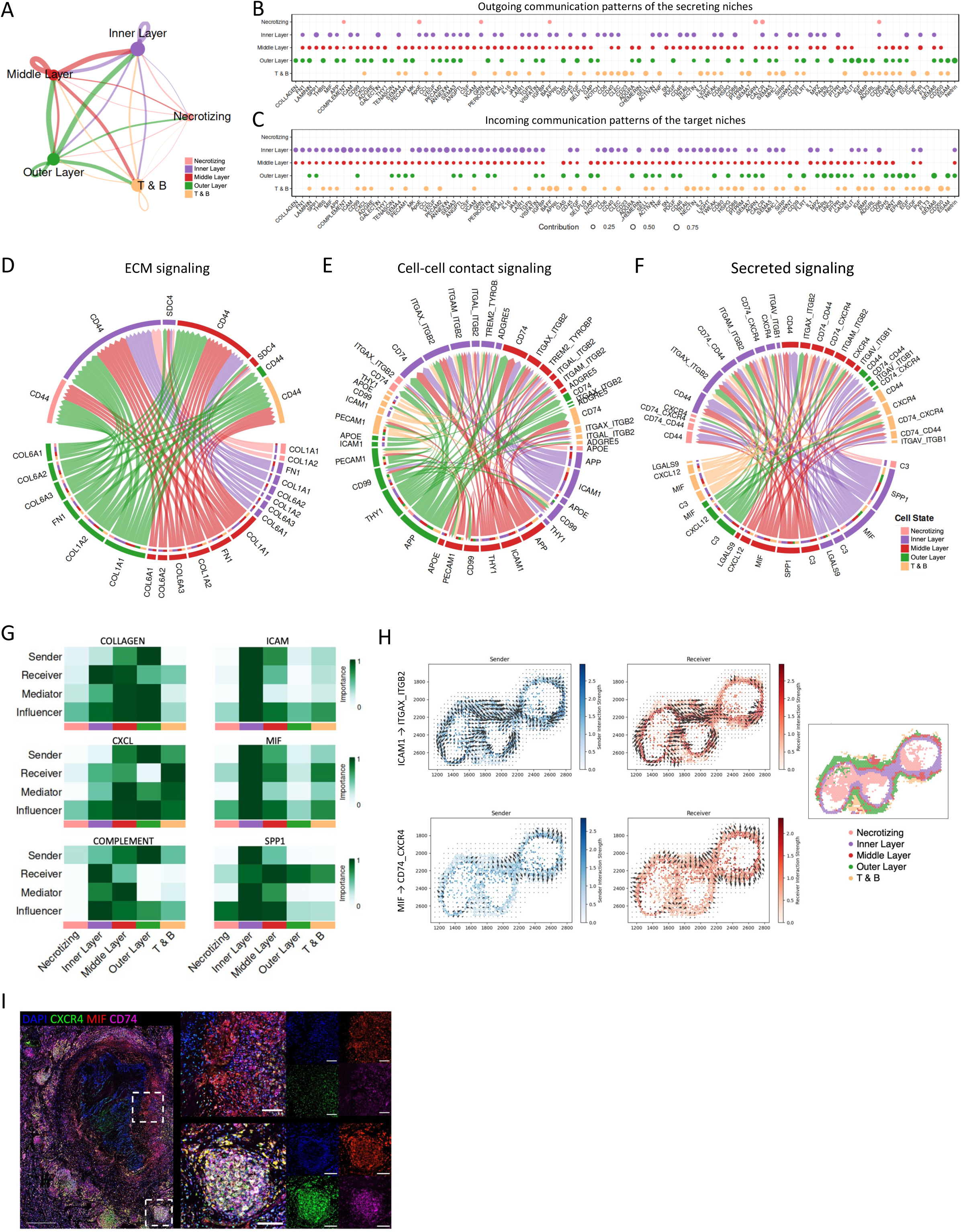
Cellular and molecular interactions across granuloma niches. (A) Network diagram showing the strength and directionality of interactions between granuloma niches. Line thickness indicates interaction strength and node size reflects the level of connectivity. (B) Outgoing and (C) incoming signal activity of each niche. (D) Circular plots depicting extracellular matrix (ECM), (E) cell-cell contact, and (F) secreted signaling interactions across granuloma niches (necrotizing core, inner niche, middle niche, outer niche, and T&B tertiary lymphoid structures). Lines represent interaction strength, with color-coded segments indicating the specific niches involved. (G) Heatmaps showing specific interaction roles for key molecules (e.g., COLLAGEN, ICAM, CXCL, MF, COMPLEMENT, SPP1) as senders, receivers, mediators, or influencers across niches. (h) Spatial maps showing sender and receiver roles for specific interactions, such as ICAM1-mediated signaling and macrophage factor MIF-CD74_CXCR4 interactions, highlighting localized signaling activity within granuloma regions. (I) Representative immunofluorescence protein staining of a MB granuloma shows the expression of MIF (red) in the inner niche and in tertiary lymphoid structures, while CXCR4 (green) and CD74 (pink) are present around the whole granuloma, especially in the tertiary lymphoid structures; Nuclei stained with DAPI (blue); Scale bar: 500 µm (large panel) and 100 µm (small panel).

To show the details of the niche-niche interaction, we demonstrate three modes of interaction: ECM interactions (Figure 3D), cell-cell contact (Figure 3E), and secreted signaling (Figure 3F). ECM interactions within MB granulomas are primarily mediated by collagens, notably COL1A1 and COL1A2. These interactions were most pronounced in the middle and outer niches, highlighting their role in maintaining granuloma structure. In contrast, cell-cell contact interactions, involving genes such as *ICAM* and *CD99*, were essential for immune cell adhesion and recruitment. Prominent secreted signaling factors included C3, MIF, CXCL12, and SPP1, further complementing immune activation, promoting chemotaxis, and bridging immune signaling with ECM remodeling.

Network centrality analysis showed prominent collagen signaling in the outer and middle niches, underscoring its importance in ECM maintenance and granuloma structure. ICAM, MIF, and SPP1 were mainly expressed in the inner niche, whereas CXCL chemokines and complement component C3 were prominent in the outer and middle niches (Figure 3G, and S3A). Interaction analysis with spatial information at individual sample levels showed the same pattern (Figure S3B).

Our analysis shed light on the mechanisms driving immune cell migration: we observed that ICAM-mediated interactions with integrins such as ITGAX and ITGB2 indicated directed movement of T cells and other immune cells toward the inner niche, facilitating stable immune engagement at the infection site (Figure 3H). Secreted factors, such as MIF and CXCL chemokines, were involved in inter-niche signaling, promoting monocytes/macrophages migration to granuloma niches (Figure 3G, 3H, and 1G). Since the sending and receiving gradients exhibit similar spatial patterns, this suggests a highly localized signaling network where MIF-producing cells are closely aligned with cells expressing its receptors. This spatial overlap also indicates an autocrine or paracrine signaling mechanism, where immune cells both secrete and respond to MIF within the same granuloma microenvironments. The presence of MIF in the inner and T&B niches was also observed on a protein level, while MIF receptors CD74 and CXCR4 were especially prominent in the T&B niche (Figure 3I). Our findings demonstrate a complex network of interactions between different granuloma niches, with distinct modes of communication driving immune cell recruitment, tissue remodeling, and overall granuloma function.

### Granuloma in MB-infected lymph nodes lacks an outer niche and shows reduced hypoxia and TGF-β signaling

During infection, mycobacterial infections can disseminate into the bloodstream, leading to extrapulmonary manifestation^43^. Apart from the lungs, lymph nodes are the second most frequently infected organs^44^. Spatial transcriptomic analysis of six explanted MB-infected lymph nodes revealed clear differences in organization and function between lymph node granuloma and pulmonary granuloma. The top 5 PCs extracted from the ST data can explain more than 50% of the variants (Figure 4A), and PC2, 4, 5, 6, and 7 are associated with the granuloma types, pulmonary and lymph node granulomas, indicating different gene expression profile between them (Figure 4b, S3A and S3E). Two morphologically lymph node granulomas could be observed: One similar to the pulmonary granuloma with a necrotizing niche surrounded by the inner and middle niches, but lacking the outer niche (n = 2; Figure 4c, red line labeled). The other type of lymph node granulomas was observed as small clusters of the necrotizing, inner, and middle niches diffusely distributed across the lymph node (n = 3; Figure 4D). As expected from their physiological structure, the T&B niche dominated the lymph nodes.

**Figure 4.**
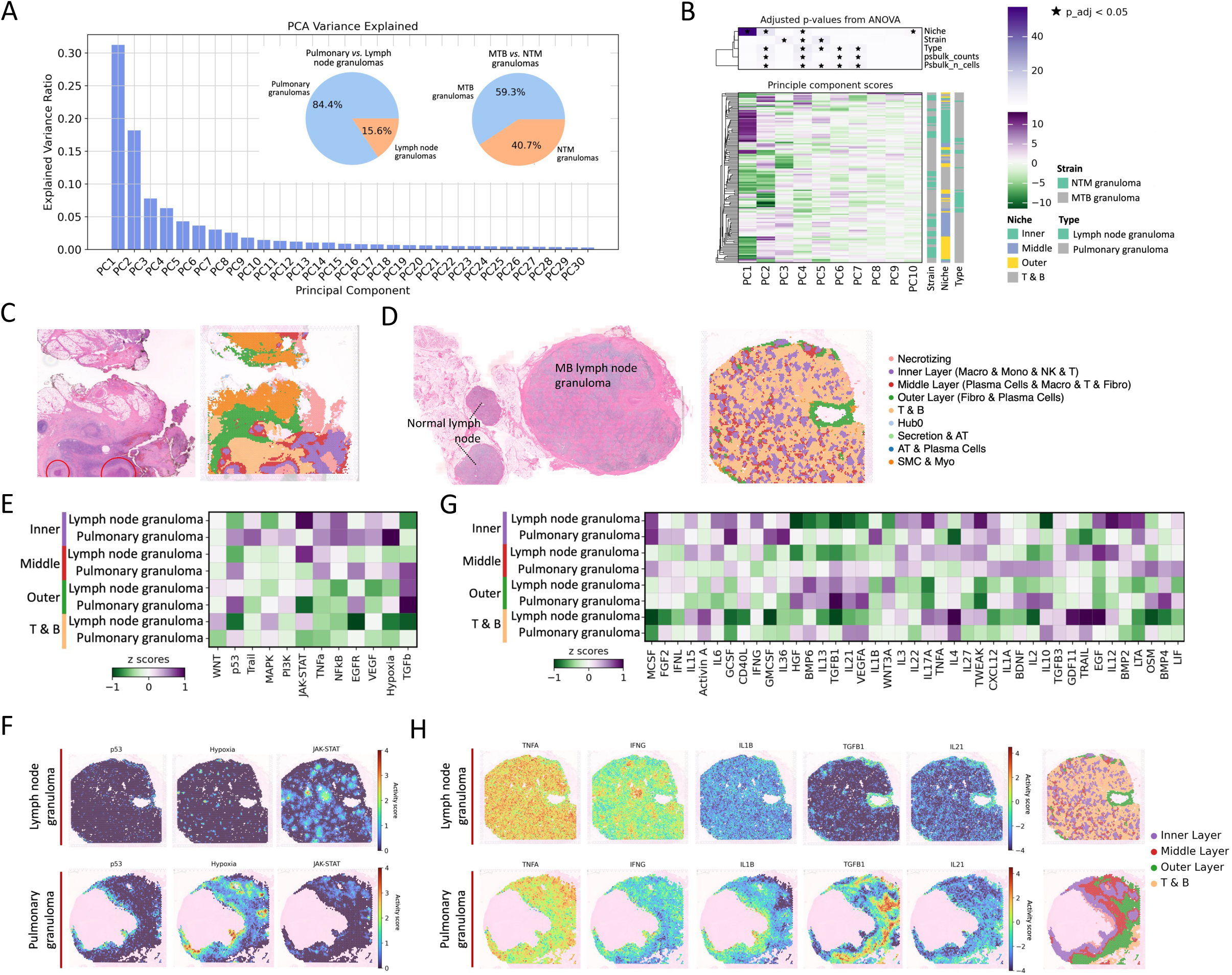
Spatial and transcriptomic comparison of pulmonary and lymph node MB granulomas. (A) PCA extracted from the ST pseudo bulk data and the sample size of pulmonary, lymph node, MTB and NTM granulomas. (B) Clustering of the PCs from niches per sample, and the correlation between PCs and spots (ST niche), strains (MTB and NTM), and types (pulmonary and lymph node). (C) Histological and (D) spatial niche annotation of pulmonary granulomas. Distinct layers include necrotic core, inner niche, middle niche, outer niche, as well as tertiary lymphoid structures (T&B niche), and additional niche regions such as secretion-associated or structural regions. (E) Heatmap showing pathway activity differences (Z-scores) across granuloma niches in pulmonary and lymph node granulomas. (F) Spatial distribution of pathway activity in pulmonary and lymph node granulomas. (G) Heatmap showing cytokine activity differences (Z-scores) across granuloma niches in pulmonary and lymph node granulomas. (H) Spatial distribution of cytokine activity in MTB lymph node granulomas (top) and pulmonary granulomas (bottom).

Granuloma-specific pathway activity also varied between pulmonary and lymph node granuloma significantly. JAK-STAT signaling was hyperactive across lymph nodes, while hypoxia and TGF-β pathways, which are critical for tissue remodeling in pulmonary granulomas, were hypoactive (Figure 4E, and 4F). Unlike the spatially confined pathway activity seen in pulmonary granulomas, JAK-STAT signaling in lymph nodes was diffuse, reflecting a more systemic immune response (Figure S4). Lymph node granulomas exhibit a pro-inflammatory Th1/Th17-driven response, with elevated TNF-α, IL-6, IL-12, IL-17A, and IFN-γ^45–47^ (Figure 4G, 4H, and S5A), promoting macrophage activation and immune cell recruitment. In contrast, lung granulomas show a fibrotic and immunoregulatory profile, enriched in BMP4, OSM, IL-13, TGF-β3, IL-21, and HGF (Figure 4G, 4H, and S5A), suggesting enhanced extracellular matrix remodeling, fibroblast activation, and immune resolution. In summary, this divergence between pulmonary and lymph node granuloma underscores the precise balance of immune signaling required for the formation of the layered niche structure that can be observed in pulmonary granuloma.

### Distinct immune signatures and hypoxia activity between MTB and NTM granulomas

To delineate the immunological and transcriptional differences between granulomas induced by MTB and NTM, we analyzed pulmonary granulomas induced by different MB strains, including 16 MTB and 11 NTM samples (Figure 4A). The spatial niches distribution revealed minimal morphological differences between MTB- and NTM-induced granulomas, with both displaying a fundamentally similar cellular architecture (Figure 5A). However, principal component analysis (PC3, 4, 5) demonstrated a strong association between granuloma transcriptional profiles and the infecting bacterial strain (Figure 4B, and S3D).

**Figure 5.**
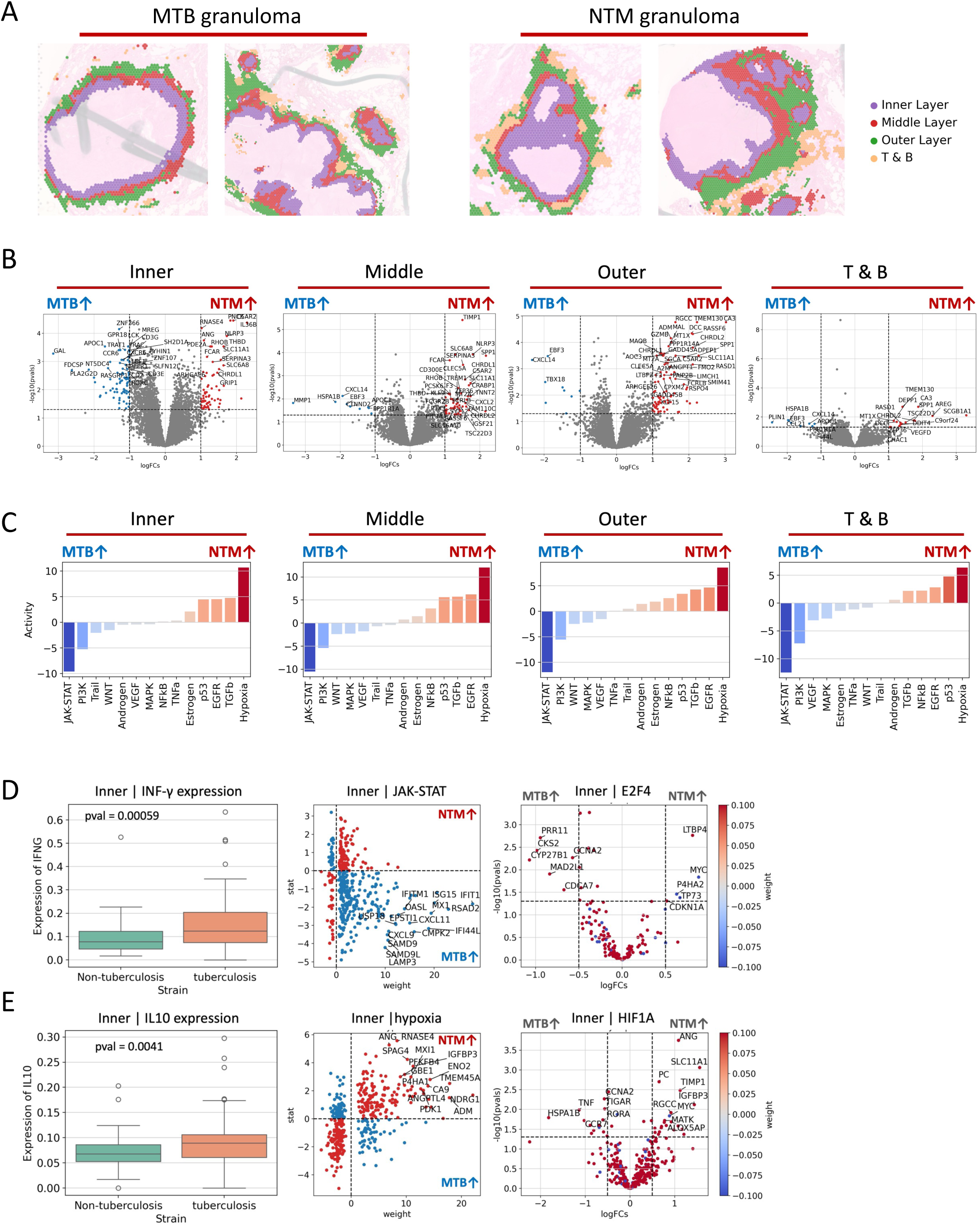
Comparative transcriptomic and pathway analysis of MTB and NTM granulomas. (A) Morphology comparison between MTB and NTM induced granulomas. (B) DEG analysis between MTB and NTM granuloma niches with pseudo-bulk data. (C) Pathway activities in MTB and NTM granuloma niches. (D) Expression of *IFN-γ*, and expression profile of genes play roles in JAK-STAT, and E2F4 activities. (E) Expression of *IL10*, and expression profile of genes play roles in hypoxia and HIF1A activities.

Across granuloma niches, MTB granulomas exhibit heightened activation of the JAK-STAT pathway with significant increased *IFN-γ* expression in inner niches (Figure 5B, 5C and 5D), promoting a Th1-driven inflammatory response. Transcription factor analysis further underscores a Th1-driven response by higher E2F4, IRF2, IRF1, and NFKB1 TF activity in the inner niche of MTB granulomas, driving monocyte differentiation, lymphocyte activation, and antigen receptor-mediated signaling^37,48^. In contrast, NTM granulomas show increased hypoxia-related activity, as indicated by differential TF activity of HIF1A (Figure 5C, 5E, and S5B), reflecting NTM lesser adaptation to hypoxia than MTB. Reduced *IL10* expression in NTM granulomas’ inner niche suggests a sustained macrophage and T cell activation, and high hypoxia reflects intense immune metabolism, bacterial control, and limited oxygen diffusion^45,49,50^ (Figure 5E). The inner niches showed functional enrichment of DEGs between MTB and NTM granulomas. The 81 upregulated genes in MTB granuloma are mainly enriched in monocytes, lymphocytes, T cell differentiation, and antigen signaling (Figure 5C). On the contrary, the 65 upregulated genes in NTM are associated with negative regulation of response to external stimulus, negative regulation of cytokine production, and response to bacterial molecules (Figure 5D). These findings suggest that both granuloma types share structural and cellular similarities, while MTB granulomas exhibit stronger immune responses and NTM granulomas are characterized by differential hypoxia signaling, emphasizing pathogen-specific immune adaptations.

### JAK-STAT signaling and tertiary lymphoid structures are hallmarks that distinguish MB from non-infection induced granulomas

To identify the transcriptional and structural features distinguishing MB granulomas from non-infection induced granulomas, we compared our 27 pulmonary MB granuloma samples with nine pulmonary sarcoidosis granuloma samples^51^. Significant structural differences were observed (Figure 6A): sarcoidosis granulomas were smaller and lacked the necrotic core and tertiary lymphoid structures – T&B niches characteristic for MB granulomas, reflecting divergent mechanisms of immune response and tissue maintenance. Despite these differences, both types of granulomas shared a similar cellular organization in their inner niches, predominantly comprising monocyte-derived macrophages, underscoring their pivotal role in granuloma formation and immune defense (Figure 6B). However, middle and outer niches were enriched of plasma cells in MB granulomas, but not in sarcoidosis granulomas (Figure 6B and 6C).

**Figure 6.**
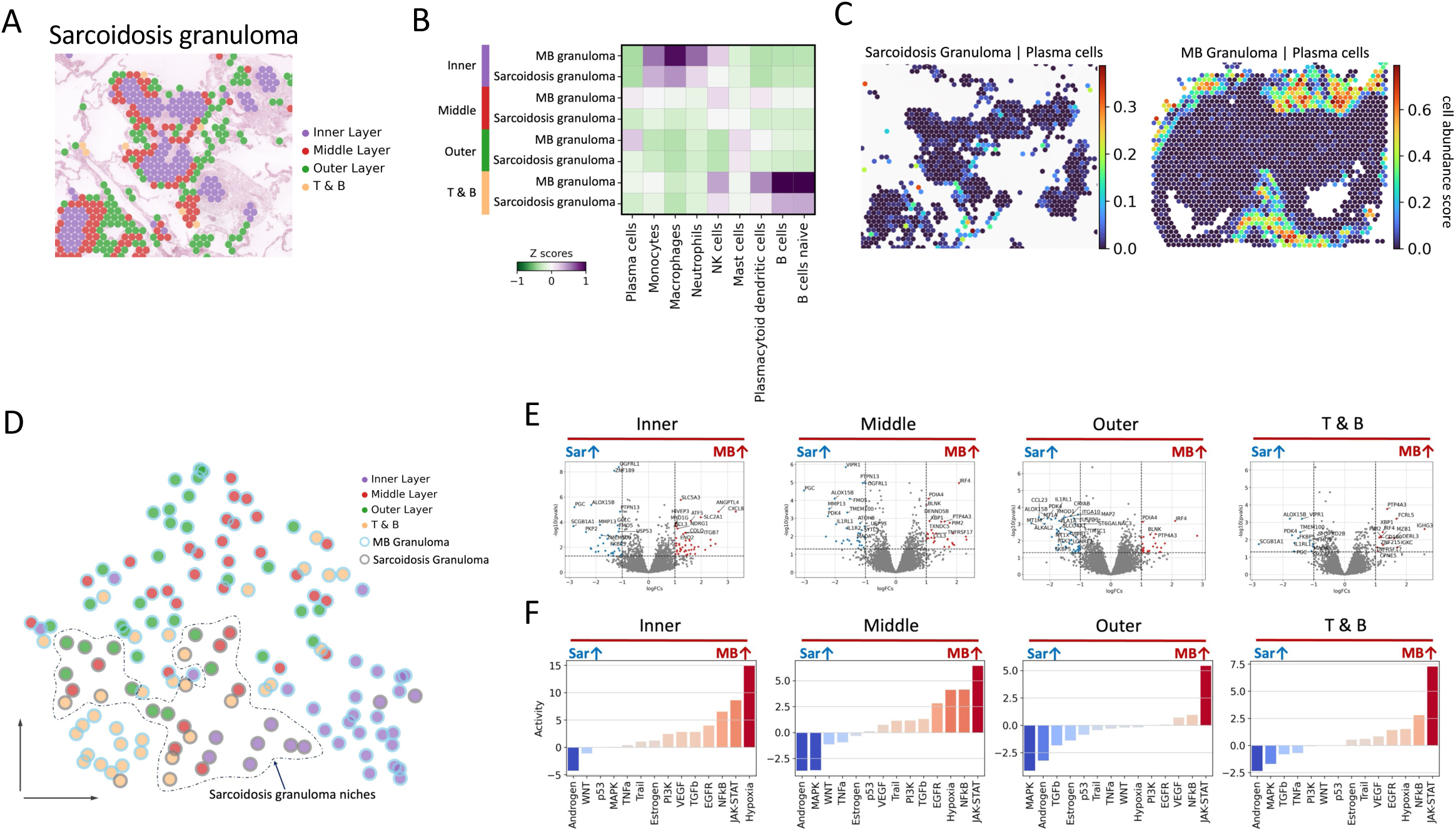
Comparative analysis of spatial niches in MB and sarcoidosis granulomas. (A) Sarcoidosis granuloma niches. Spatial niches were identified and annotated. (B) Heatmap showing cell-type association with each granuloma niche across both MB and sarcoidosis samples. (C) Plasma cells abundance in Sarcoidosis and MB granulomas. The abundance is presented by the MLM score based on the plasma cell marker genes’ expression. (D) UMAP visualization of spatial niches highlighting the difference in clustering patterns between MB and sarcoidosis granulomas. (E) Differentially expressed genes (DEGs) between matched niches from MB and sarcoidosis granulomas. Each plot represents a comparison of corresponding niches, indicating gene upregulation and downregulation between the granuloma types. (F) Inferred pathway activity scores for the matched niches of MB versus sarcoidosis granulomas. Sar stands for sarcoidosis.

Pseudo-bulk analysis revealed transcriptional divergence based on granuloma etiology. Sarcoidosis granulomas showed tighter clustering of spatial gene expression spots in the UMAP, indicating greater homogeneity within lesions compared to the heterogeneous spatial profiles observed in MB granulomas (Figure 6D). This variability in MB granulomas likely reflects the complex and dynamic immune responses required to combat a persistent bacterial infection.

To better understand these differences, we examined gene expression and pathway activities across comparable niches. Differential gene expression analysis highlighted the upregulation of *BLNK* and *TNFRSF17* in MB granulomas (Figure 6E, Middle, Outer, and T&B niches), genes associated with B cell development. These findings underscore the more pronounced adaptive immune responses in MB granulomas, particularly through plasma and B cell involvement. The inner and middle niches of the MB granuloma exhibited higher hypoxia signaling, possibly reflecting a low-oxygen microenvironment, in addition to higher JAK-STAT signaling across all common granuloma niches (Figure 6F, S6A, S6B, S6C, and S6D). In contrast, sarcoidosis granulomas showed increased MAPK signaling in the middle and outer niches (Figure 6F), suggesting distinct mechanisms of tissue remodeling and immune cell activation.

## Discussion

In this study, we provide the first spatial transcriptomic atlas of tuberculosis and non-tuberculous mycobacterial disease. MB-induced granulomas reveal a highly organized immune microenvironment coordinating pathogen containment and tissue remodeling. We identified five distinct niches: (i) the necrotizing niche marked by extensive cell death and low transcripts; (ii) the inner niche, rich in macrophages, JAK-STAT activity, and hypoxia signaling, indicating active immune responses; (iii) the middle niche, a transitional zone integrating inflammation and ECM remodeling; (iv) the outer niche, dominated by fibroblasts and crucial for ECM remodeling and structural stability; and (v) T&B niches, which reflect tertiary lymphoid structures, supporting antigen presentation and adaptive immunity. A spatial gradient from immune activation at the core to ECM remodeling at the periphery was evident, consistent with ligand-receptor interactions. Findings were replicated in an independent cohort (Figure S7).

As there is no “healthy” or “normal” analog to MB granulomas, we sought to study basic principles of MB granuloma architecture and homeostasis by comparing different affected organs (lung vs. lymph nodes), different inducing strains (TB vs. NTM) and by comparing them to non-infectious granulomas of sarcoidosis patients. Granuloma form tuberculosis differed from non-tuberculous MB disease: MTB granulomas displayed strong IFN-γ and JAK-STAT activation, whereas NTM granulomas exhibited heightened hypoxia and immunoregulatory signaling, reflecting distinct host response across different granuloma types. Both the host or the pathogen could contribute to this: On the one hand, the observed distinct immune landscapes of MTB and NTM granulomas suggest that different mycobacterial species could provoke adapted host responses to optimize their survival. MTB, as an obligate human pathogen, can e.g. improve its survival in the host through promoting JAK-STAT-induced apoptosis of macrophages^52^. In contrast, the upregulation of hypoxia-related pathways and reduced JAK-STAT signaling in NTM granulomas suggests that NTM profit from a naturally constrained immune response in a hypoxic environment rather than modulating or disabling the host response. On the other hand, NTM-infections primarily affects the immunocompromised host or host with preexisting pulmonary disease, lacking adequate anti-MB immune response. E.g., it is well known from genetic studies, that the IFN-γ axis is essential for protection against MB, and that its absence can lead to severe, disseminated MB infection due to failure of bacterial containment^53^. Evaluation of lymph node granulomas revealed a pro-inflammatory Th1/Th17-driven response, but lacked a dedicated hypoxia central niche and an outer fibroblast-rich niche. The absence of an outer niche is consistent with findings that describe lymph node granulomas as coalescing structures that displace the normal lymph node architecture^54,55^. Structural support may not be essential for lymph node granuloma integrity due to the inherent density of the lymph node. This is in contrast to the thin-walled alveoli of aerated lungs, where the formation of an outer granuloma niche in lung granulomas may be a prerequisite for MB containment. Comparison with sarcoidosis granulomas as a disease control revealed the following two important observations: First, *SPP1* and *CHI3L1*, both involved in inflammation, tissue remodeling and M2 macrophage polarization, were highly expressed by macrophages in both sarcoidosis and MB granulomas, implicating a general role in granuloma formation^51,56,57^. Second, the lack of plasma cells and tertiary lymphoid structures around sarcoidosis granuloma affirms their importance in MB granuloma while implicating that they are not necessary for granuloma formation itself. Nonetheless, B cells of the granuloma-associated lymphoid tissue facilitate MB phagocytosis by secretion of MB-specific IgM antibodies^58^, thereby promoting immunity. Whether they confer risk for autoimmune disease in TB patients, is a matter of same debate, as tertiary lymphoid structure can become dysregulated over time^59^. Taken together, our comparative analysis between granulomas of different MB strains, affected organs and sarcoidosis as disease control revealed shared and unique features of granulomatous inflammation, highlighting disease-specific mechanisms.

Post-tuberculosis lung disease is a frequently progressive fibrotic sequela of tuberculosis that persists even after the infection has been cured^60^. Our study suggests that the relevant cellular foundations for post-tuberculosis lung disease are already laid during active MB infection: Fibroblasts of the outer wall are an integral constituent of granuloma architecture. Coinciding with high TGF-β activity, they express a multitude of collagens, as well as CTHRC1, recently described as hallmark gene of myofibroblasts in the IPF lung^28^. Those fibroblasts acquired the inflammatory phenotype and signal especially through the CXCL12-CXCR4 axis to the immune cells in middle and inner layer^28,61^. Last, also profibrotic macrophages were observed in the inner and middle wall, with prominent MIF and SPP1 signaling through e.g. receptors CD44 and CD74^29,62^. Therapeutically, it is crucial to identify the temporal window during which fibroblasts are no longer essential for mycobacterial containment, thereby enabling the safe implementation of anti-fibrotic interventions to mitigate post-tuberculosis lung disease.

Adding to the therapeutic perspective, our findings suggest that precision host-directed therapies should be tailored to target granuloma-specific niches. For example, enhancing immune activation in the inner niches in NTM disease, where macrophages and T cells interact with MB, could be achieved by boosting IFN-γ and TNF-α signaling while counteracting T cell exhaustion via PD-1/PD-L1 blockade^63^ or IDO1 inhibition^64^. While the benefits of IFN-γ in active tuberculosis still remain inconclusive, partially due to the lack of large clinical trials^65^, additionally targeting MIF, a key regulator of macrophage inflammatory function, may enhance MB-reactive immune responses. As another example, patients with fibrotic granulomas may benefit from anti-fibrotic therapies such as nintedanib^66^ and pirfenidone^67^ to minimize the loss of functional lung tissue due to fibrosis. Future studies integrating multi-omics analyses with *in vivo* imaging-based diagnostics will be essential for characterizing individual granuloma phenotypes and guiding precision host-directed therapies.

Our study is not without limitations. First, the resolution (100µm) of the utilized 10x Genomics Visium platform lacks single-cell resolution and is restricted to a transcriptomic analysis of “only” ∼19,000 genes. Future studies using single-cell resolution will provide a more comprehensive view of granuloma biology. Second, the probe-based Visium platform is unable to detect mycobacterial RNA, so interactions between the pathogen and the host could not be explored. Third, in the NTM subset of this study, we cannot resolve whether observed differences between TB and NTM granuloma are due to differences in the MB strains or the susceptible host. Further studies are therefore required to tease apart host and the pathogen-effect of the observed substantial spatial transcriptomics differences between TB and NTM granuloma.

In conclusion, our findings refine the understanding of the spatial architecture, homeostasis and immune regulation in MB granulomas, emphasizing their active, dynamic nature rather than passive immune aggregates. The layered immune organization, metabolic specialization, pathogen-specific adaptations, and mesenchymal containment suggest that MB treatment strategies must consider granuloma microenvironments as distinct functional units. Last, we hope that our open-access and easily accessible resource https://lab-li.ciim-hannover.de/mb-granuloma/ will advance MB research, paving the way towards precision therapies for patients suffering from tuberculosis or NTM disease.

## Resource availability

Our raw data were deposited in EGA database as FASTQ format (EGAS50000000998). The processed data is presented in our webtool https://lab-li.ciim-hannover.de/mb-granuloma/. All codes used for the analysis in this paper is uploaded to: https://github.com/CiiM-Bioinformatics-group/MB_Granuloma/.

## Ethics statement

The study was approved by the institutional ethics committees (MHH: IRB # 10142_BO_K_2022; Achen: EK-23350; Yale: 2000036632)

## Acknowledgments

This work is supported an ERC Starting Grant (948207, ModVaccine) to Y.L., by the Ministry of Science and Culture of Lower Saxony through funds from the program Zukunft. Niedersachsen of the Volkswagen Foundation for the ‘CAIMed – Lower Saxony Center for Artificial Intelligence and Causal Methods in Medicine’ project (grant no. ZN4257) to Y.L., the Deutsche Forschungsgemeinschaft (DFG, German Research Foundation) under Germany’s Excellence Strategy - EXC 2155 - project number 390874280 to Y.L., This project is supported by the Lower Saxony MWK Sprung Fund to C.X. (19777006). J.C.S. is supported by the Fritz Thyssen Foundation (10.21.2.021MN) and the Else Kröner-Fresenius Foundation (2023_EKCS.18). L.N. and J.C.S. are supported by the Ann Theodore Foundation and the German Center for Lung Research (FKZ 82DZL002B1, FKZ 82DZL002C1 & FKZ 82DZLT82C1). MK is supported by the Deutsche Forschungsgemeinschaft (KA 6005/1-1).

## Author contributions

X.J. and L.C. performed the experiments, analyzed the data, and wrote the manuscript. L.Z., A.A., and X.Z. contributed to experimental work. Y.X. and N.v.U. developed the interactive web tool for processed data visualization. Y.L., J.C.S., C.-J.X., F.R., R.H., and D.D.J. supervised the study, provided critical suggestions, and assisted with manuscript writing. L.N., J.D., J.C.K., M.K., J.H., H.S., M.M.H., N.K., T.W., and J.F. contributed to the provision and preparation of FFPE tissue samples. All authors read and approved the final version of the manuscript.

## Declaration of interests

J.F. has received fees for consultations from AstraZeneca unrelated to the submitted work. H.M. in MMH has received fees for consultations or lectures from 35Pharma, Acceleron, Actelion, Aerovate, AOP Health, Bayer, Ferrer, Gossamer, Inhibikase, Janssen, Keros, MSD and Novartis. F.C.R. reports grants or contracts from German Center for Lung Research (DZL), German Center for Infection Research (DZIF), IMI (EU/EFPIA) and iABC Consortium, Mukoviszidose Institute, Novartis, and Insmed; consulting fees or participation on a advisory board from Boehringer Ingelheim, Parion Sciences, Chiesi, Sanofi, and Insmed; payment or honoraria for lectures, presentations, speakers fees, or educational events from I!DE Werbeagentur, AstraZeneca, Insmed, Grifols, Universitätsklinikum Hamburg-Eppendorf, and Cliniqo; support for attending meetings and/or travel from German Kartagener Syndrome and Primary Ciliary Dyskinesia Patient Advocacy Group and German Cystic Fibrosis Patient Advocacy Group; he received fees for clinical trial participation paid to his institution by AstraZeneca, Boehringer Ingelheim, Insmed, Novartis, Parion Sciences, University of Dundee, Vertex and Zambon.

## Supplemental information titles and legends

**Figure S1** (A) Detected genes, counts and the genes per Spot after quality control. (B) NMFs extracted from the ST data and plotted in 2D for an example sample.

**Figure S2** (A) Niche-specific DEGs present in over 50% of spots in each niche, but in fewer than 30% of other niches, identifying the most specific marker genes of each granuloma niche. (B) Comparison between inner and middle, middle and outer niches. (C) Maker genes for Trail, Hypoxia, JAK-STAT, and TGFb pathway in 2D for an example sample. (D) Biological process enrichment results of the DEGs showed in Figure 2D.

**Figure S3** (A) Ranking of the Ligand-receptor interaction in MB granuloma niches. (B) Niche-niche interaction within an individual sample with spatial information. (C) (D) PCA plot showing the difference in gene expression across MB granuloma types. (E) Comparison between pulmonary and lymph node granulomas across niches.

**Figure S4** Biological process enrichment analysis of the DEGs in Figure 4F in each niche.

**Figure S5** (A) Cytokine activity difference between pulmonary and lymph node granuloma in whole granuloma. (B) TF activity difference between NTM and MTB. (C) Biological process enrichment of the 81 upregulated genes in MTB granulomas, and (D) the 65 upregulated genes in NTM granulomas.

**Figure S6** The DEGs make contribution to the differential pathway activity between Lymph node and Pulmonary granulomas. (A) Expression level of maker genes for Hypoxia, and JAK-STAT pathway in the inner niche. (B) Expression level of maker genes for JAK-STAT pathway in the middle niche. (C) Expression level of maker genes for JAK-STAT pathway in the outer niche. (D) Expression level of maker genes for JAK-STAT pathway in the T&B niche. Red: genes make positive contribution to pathway activity in lymph node granulomas. Blue: genes make positive contribution to pathway activity in pulmonary granulomas.

**Figure S7** (A) Genes per spot and counts detected in the replication cohort. (B) Replication of the JAK-STAT, hypoxia, and TGFb pathway activity. (C) Cytokine activity replication.

**Table S1** Sample information containing sex, age, and infected strains.

**Table S2** Summary of the original spaceranger outputs information for all the samples used in this study.

**Table S3** Antibodies and their dilution used in this study

## STAR Methods

### Spatial transcriptomic profiling of the MB granuloma samples

Spatial transcriptomics was performed using the 10x Genomics Visium Spatial Gene Expression platform, which utilizes an 11mm x 11mm probe array to capture spatial gene expression information for the 32 samples in the discovery cohort, 6.5mm x 6.5mm probe array for the replication cohort. Biobanked FFPE lung tissue samples containing mycobacteria-associated granulomatous lesions of patients providing informed consent were collected from three different institutions: Hannover Medical School (Germany), University Hospital RWTH Aachen (Germany), and Yale School of Medicine (USA). Samples in the replication cohort are all from the Hannover Medical School. Before spatial analysis, H&E staining was carried out to assess tissue morphology and select regions of interest.

### Slide preparation and spatial profiling

Spatial transcriptomics was carried out according to the manufacturer’s instructions (10x Genomics, CG000408, CG000409, CG000407). In short: tissue sections were H&E stained, imaged, destained, and decrosslinked before on-slide probe hybridization. After probe ligation, the CytAssist was used to transfer probes onto the Visium slides. Probes were extended to generate spatially barcoded cDNA. This spatially barcoded cDNA was then amplified and used for library construction. The constructed libraries were sequenced using the Illumina sequencing platform to generate spatial gene expression data. The RNA probe hybridization mix was prepared using the Visium FFPE Human Transcriptome Probe Set (10x Genomics, Visium Human Transcriptome Probe Set v1.0), following the manufacturer’s recommended dilution and conditions. In situ extension reactions were then carried out to generate spatially cDNA. Following cDNA synthesis, spatially barcoded cDNA was amplified via PCR, purified, and used for library construction using the Visium Spatial Gene Expression library preparation protocol for FFPE samples. Final libraries were quantified, quality-controlled using Bioanalyzer, and stored at −20 °C prior to sequencing.

### cDNA Sequencing and Data Processing

Sequencing was performed on an Illumina NovaSeq 6000 platform using paired-end 50 bp reads (PE50). This sequencing configuration ensured sufficient read depth and quality for spatial transcriptomic analysis of FFPE-derived samples. The raw base call (BCL) files from the NovaSeq runs were converted to FASTQ format using the spaceranger mkfastq command from Space Ranger v3.0 (10x Genomics). Subsequently, gene expression data was generated by aligning reads to the appropriate reference transcriptome (refdata-gex-GRCh38-2020-A) using the Space Ranger v3.0 count pipeline, assign unique molecular identifiers (UMIs), and generate count matrices representing gene expression across spatial spots on the tissue. This process produced spatially resolved gene count matrices for downstream analysis.

### Cell type deconvolution

Cell type deconvolution was performed using the Cell2location package^23^ (version 0.1.3), which allows for accurate inference of cell type proportions at each spatial spot. An average cell count per spot was estimated at 20, providing a high-resolution view of the cellular landscape within each granuloma niche. Cell2location utilized reference single-cell RNA sequencing data (HLCA v2^25^, anno4 was used) to assign cell types and quantify their abundance across the spatial transcriptomics data, aiding in the detailed characterization of the cellular architecture in granulomas.

### Sample integration and harmonization

The gene count matrices from cohorts were integrated and harmonized separately using RAPIDS-SingleCell^68^ (version 0.10.5) and Scanpy^69^ (version 1.10.2). RAPIDS-SingleCell was employed to efficiently align the high-dimensional data, leveraging GPU acceleration for faster computation, while Scanpy provided the necessary preprocessing, normalization (10^4^), data quality control (count > 2000) and batch correction steps. The integrated data was saved as a h5ad file for subsequent analysis.

### Spatial niches clustering and annotation

Leiden algorithm clustering^70^ was used to identify distinct spatial niches by analyzing the spots from all samples. The clustering was informed by deconvoluted cell types derived from the Cell2location tool, alongside known marker genes for lung tissue cell types. Each spot was annotated based on its dominant cell type composition.

### Pathway and cytokine activity analysis

Pathway activity analysis was conducted using the Decoupler package^71^ (version 1.7.0), with pathway scoring performed based on the Progeny database^72^. The multivariate linear model (MLM) was applied to map pathway activity scores to the spatial spots, allowing for a detailed spatial understanding of pathway activity across the tissue. For cytokine activity analysis, the CellTypist database^30,31^ was used to determine cytokine activity at each spot. Univariate linear modeling (ULM) was utilized to assign cytokine activity scores, providing insight into localized immune signaling patterns. Pathway enrichment analysis was performed using the KEGG^73^ or Reactome^74^ pathway database. Over-representation analysis (ORA) was used to calculate enrichment scores.

### Niche-niche interaction analysis

Niche-nich interaction analysis was conducted using the CellChat^75^ package (version 2.1.2) on all integrated samples, with default settings. For individual sample analysis, ECM and cell-cell contact ligand-receptor (LR) pairs were analyzed with contact dependent set to TRUE, while secreted LR pairs were analyzed with interaction range set to 250. The Commot^76^ package (version 0.0.3) was used to analyze the gradient trends of LR interaction activity. Additionally, the ConnectomeDB2020 LR pair database^75,76^ was used via the stlearn^77^ package (version 0.4.12) for similar analyses. The results obtained from CellChat and ConnectomeDB2020 were consistent across all samples.

### pseudo-bulk RNAseq analysis

pseudo-bulk RNA sequencing analysis was performed using the Decoupler^71^ (version 1.7.0). Gene counts from spots annotated in the same clusters were summed. Normalization and log(x+1) transformation were applied to the aggregated gene counts to prepare the data for downstream analyses.

### DEGs and pathway enrichment analysis

Comparative analysis was performed using DESeq2 approach based on the pseudo-bulk RNAseq data with the Decoupler package^71^ (version 1.7.0), comparing different spatial niches or the same niches between MTB and sarcoidosis granulomas. The spatial transcriptomics data of pulmonary sarcoidosis granuloma was employed with author’s permission (GEO: GSE288607). The functional enrichment analysis of DEGs was performed with the clusterProfiler R package^78^ (version 4.6.2).

### Immunofluorescence protein staining

The slides were rehydrated using a series of xylene (Carl Roth, # CN80.1) and ethanol (Otto Fischar, # 27690). Tissue was then decrosslinked by incubation in 95°C water with 1% Antigen Unmasking Solution (Vector Laboratories, # H3301) for 20 minutes. Afterward, the slides were cooled to room temperature in 1X PBS (Chemsolute, # 8418) for 20 minutes. Blocking of nonspecific antibody binding was reduced by incubation with 2.5% v/v normal donkey serum (Biozol, # JIM-017-000-121). The slides were then incubated with the primary antibodies for 1 h. After rinsing in 1X PBS (2 min, twice), secondary antibodies were applied for 1 h in the dark. Slides were rinsed in 1X PBS (2 min, twice) and incubated with pre-conjugated antibodies for 1 h. To minimize background, the Vector TrueView Autofluorescence Quenching Kit (Vector Laboratories, # SP-8400-15) was applied as per the manufacturer’s instructions. After rinsing in 1X PBS (2 min, twice), the slides were mounted with DAPI-containing Antifade Mounting Medium (Vector Laboratories, # H-1800-10) and coverslippe. Slides were stored in the dark at 4°C until imaging.

For imaging, a Zeiss Axio Scan 7 equipped with a 20x Plan Neofluar Objective was used. For emission filters, the 96 HE BFP 450/40,38 eGFP 525/50, 43 HE DsRed 605/70, and 50 Cy5 690/50 broad pass filters were used. Fluorescence intensity was adjusted for each slide individually, aiming at a maximum of 5% oversaturation. Details of antibodies used, including clones, concentrations, and suppliers, are listed in Table S3.

## Notes

### Competing Interest Statement

The authors have declared no competing interest.

