## Supplemental Figures for "A multicenter spatial transcriptomics atlas of human tuberculosis and non-tuberculous mycobacterial disease"

Figure S1

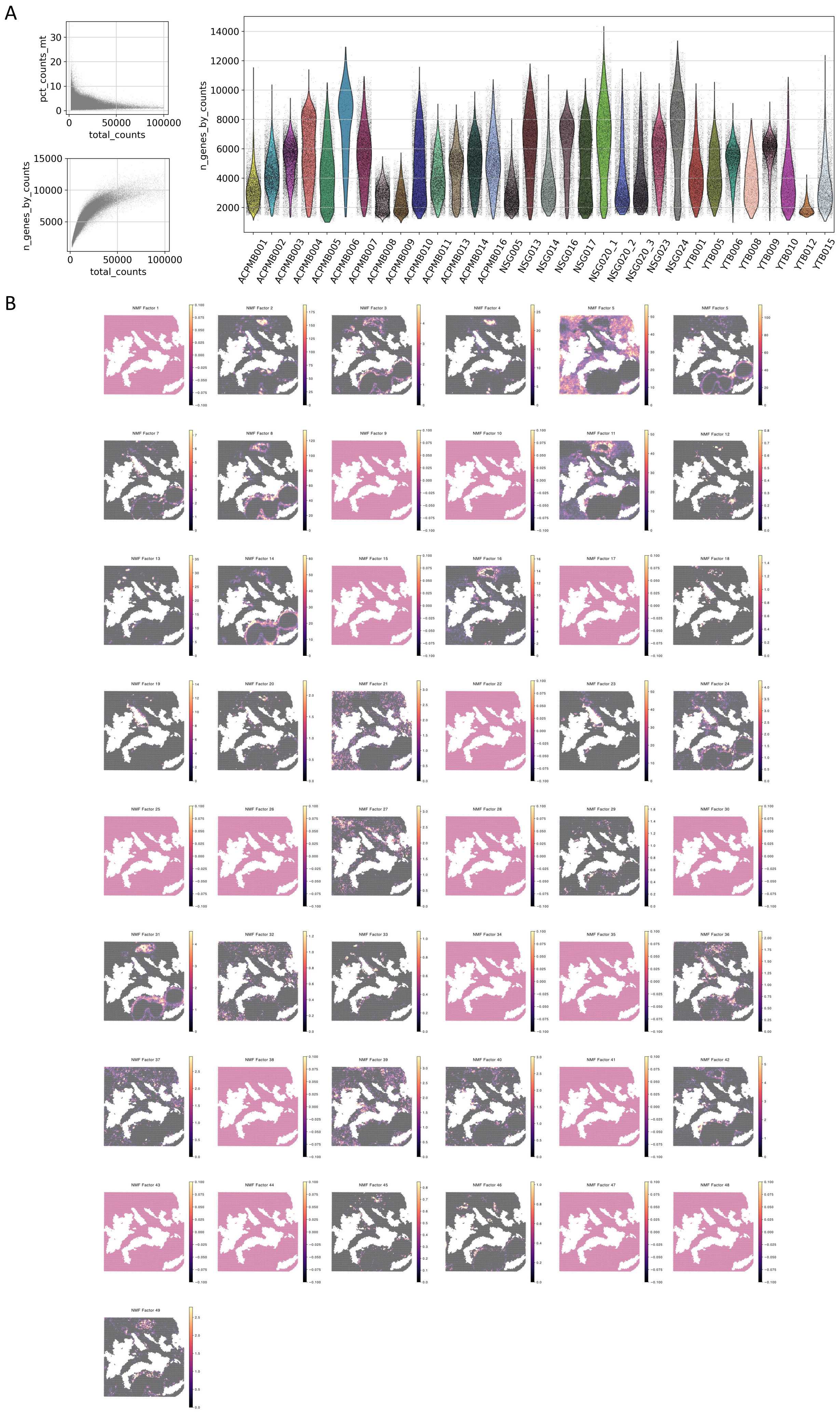
A
pct_counts_mt
total_counts
n_genes_by_counts
ACPMB001
ACPMB002
ACPMB003
ACPMB004
ACPMB005
ACPMB006
ACPMB007
ACPMB008
ACPMB009
ACPMB010
ACPMB011
ACPMB013
ACPMB014
ACPMB016
NSG005
NSG013
NSG014
NSG016
NSG017
NSG020_1
NSG020_2
NSG020_3
NSG023
NSG024
YTB001
YTB005
YTB006
YTB008
YTB009
YTB010
YTB012
YTB015
B

Figure S2

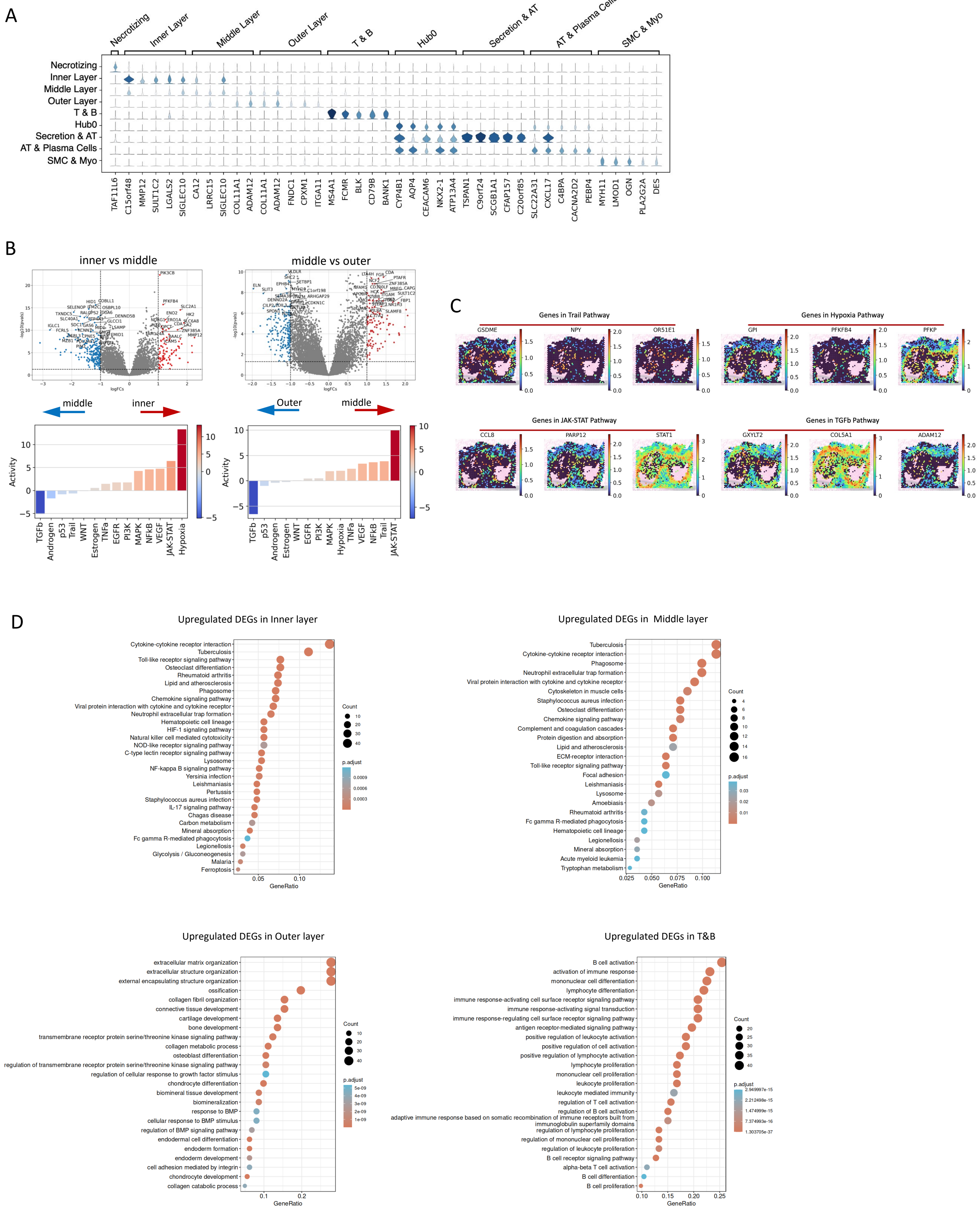
A
Necrotizing
Inner Layer
Middle Layer
Outer Layer
T & B
Hub0
Secretion & AT
AT & Plasma Cells
SMC & Myo
TAF11L6
C15orf48
MMP12
SULT1C2
LGALS2
SIGLEC10
CA12
LRRC15
SIGLEC10
COL11A1
ADAM12
COL11A1
ADAM12
FNDC1
CPXM1
ITGA11
MS4A1
FCMR
BLK
CD79B
BANK1
CYP4B1
AQP4
CEACAM6
NKX2-1
ATP13A4
TSPAN1
C9orf24
SCGB1A1
CFAP157
C20orf85
SLC22A31
CXCL17
C4BPA
CACNA2D2
PEBP4
MYH11
LMOD1
OGN
PLA2G2A
DES
B
inner vs middle
middle vs outer
logFCs
-log10(pvals)
middle
inner
Outer
middle
Activity
TGFb
Androgen
p53
Trail
WNT
Estrogen
TNFa
EGFR
PI3K
MAPK
NFkB
VEGF
JAK-STAT
Hypoxia
TGFb
p53
Androgen
Estrogen
WNT
EGFR
PI3K
MAPK
Hypoxia
TNFa
VEGF
NFkB
Trail
JAK-STAT
C
Genes in Trail Pathway
GSDME
NPY
OR51E1
Genes in Hypoxia Pathway
GPI
PFKFB4
PFKP
Genes in JAK-STAT Pathway
CCL8
PARP12
STAT1
Genes in TGFb Pathway
GXYLT2
COL5A1
ADAM12
D
Upregulated DEGs in Inner layer
Cytokine-cytokine receptor interaction
Tuberculosis
Toll-like receptor signaling pathway
Osteoclast differentiation
Rheumatoid arthritis
Lipid and atherosclerosis
Phagosome
Chemokine signaling pathway
Viral protein interaction with cytokine and cytokine receptor
Neutrophil extracellular trap formation
Hematopoietic cell lineage
HIF-1 signaling pathway
Natural killer cell mediated cytotoxicity
NOD-like receptor signaling pathway
C-type lectin receptor signaling pathway
Lysosome
NF-kappa B signaling pathway
Yersinia infection
Leishmaniasis
Pertussis
Staphylococcus aureus infection
IL-17 signaling pathway
Chagas disease
Carbon metabolism
Mineral absorption
Fc gamma R-mediated phagocytosis
Legionellosis
Glycolysis / Gluconeogenesis
Malaria
Ferroptosis
GeneRatio
Count
p.adjust
Upregulated DEGs in Middle layer
Tuberculosis
Cytokine-cytokine receptor interaction
Phagosome
Neutrophil extracellular trap formation
Viral protein interaction with cytokine and cytokine receptor
Cytoskeleton in muscle cells
Staphylococcus aureus infection
Osteoclast differentiation
Chemokine signaling pathway
Complement and coagulation cascades
Protein digestion and absorption
Lipid and atherosclerosis
ECM-receptor interaction
Toll-like receptor signaling pathway
Focal adhesion
Leishmaniasis
Lysosome
Amoebiasis
Rheumatoid arthritis
Fc gamma R-mediated phagocytosis
Hematopoietic cell lineage
Legionellosis
Mineral absorption
Acute myeloid leukemia
Tryptophan metabolism
GeneRatio
Count
p.adjust
Upregulated DEGs in Outer layer
extracellular matrix organization
extracellular structure organization
external encapsulating structure organization
ossification
collagen fibril organization
connective tissue development
cartilage development
bone development
transmembrane receptor protein serine/threonine kinase signaling pathway
collagen metabolic process
osteoblast differentiation
regulation of transmembrane receptor protein serine/threonine kinase signaling pathway
regulation of cellular response to growth factor stimulus
chondrocyte differentiation
biomineral tissue development
biomineralization
response to BMP
cellular response to BMP stimulus
regulation of BMP signaling pathway
endodermal cell differentiation
endoderm formation
endoderm development
cell adhesion mediated by integrin
chondrocyte development
collagen catabolic process
GeneRatio
Count
p.adjust
Upregulated DEGs in T&B
B cell activation
activation of immune response
mononuclear cell differentiation
lymphocyte differentiation
immune response-activating cell surface receptor signaling pathway
immune response-activating signal transduction
immune response-regulating cell surface receptor signaling pathway
antigen receptor-mediated signaling pathway
positive regulation of leukocyte activation
positive regulation of cell activation
positive regulation of lymphocyte activation
lymphocyte proliferation
mononuclear cell proliferation
leukocyte proliferation
leukocyte mediated immunity
regulation of T cell activation
regulation of B cell activation
adaptive immune response based on somatic recombination of immune receptors built from immunoglobulin superfamily domains
regulation of lymphocyte proliferation
regulation of mononuclear cell proliferation
regulation of leukocyte proliferation
B cell receptor signaling pathway
alpha-beta T cell activation
B cell differentiation
B cell proliferation
GeneRatio
Count
p.adjust

Figure S3

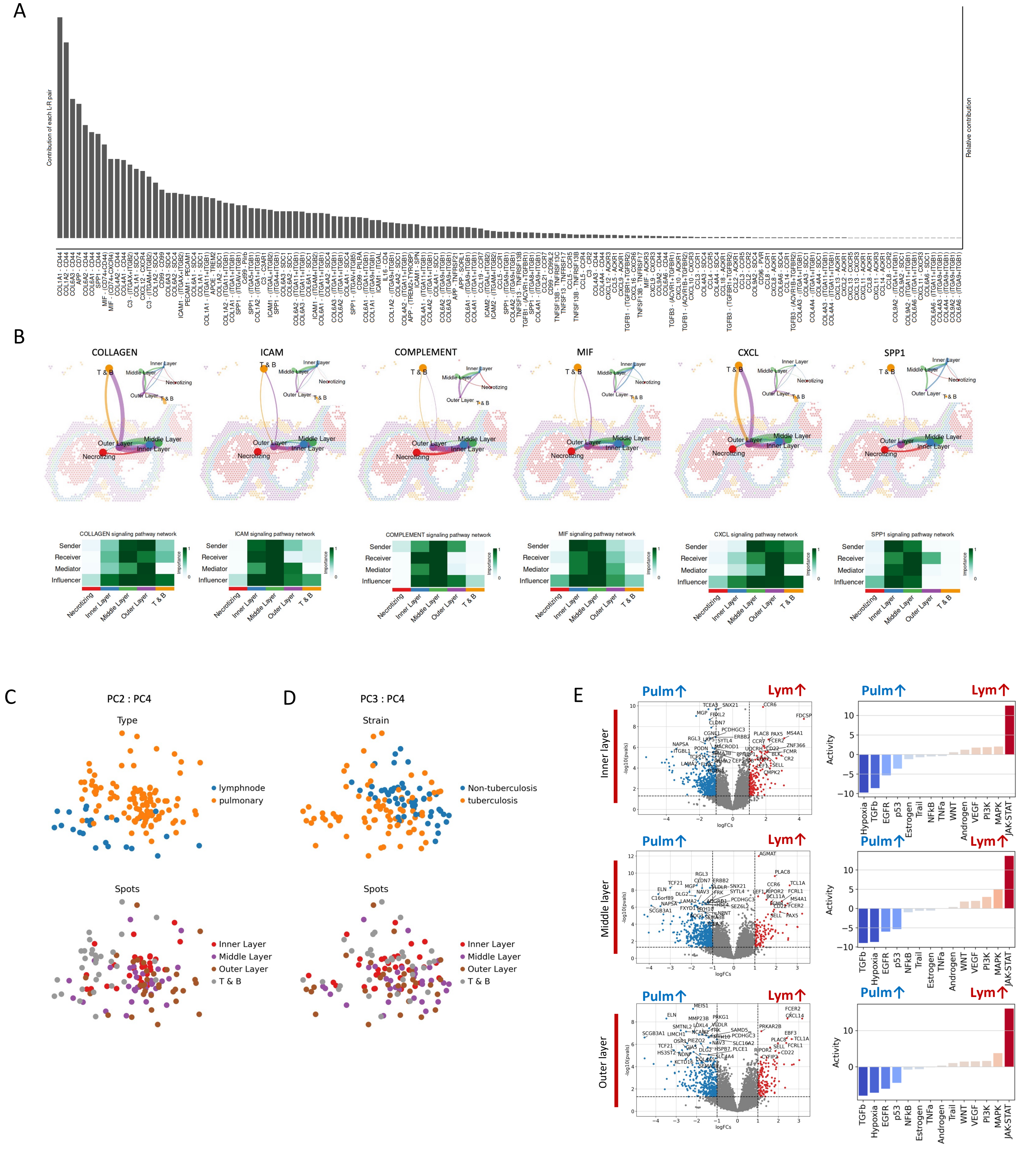


Supplementary Fig. 3

(a) Ranking of the Ligand-receptor interaction in MB granuloma niches. (b) Niche-niche interaction within an individual sample with spatial information . (c) (d) PCA plot showing the difference in gene expression across MB granuloma types. (e) Comparison between pulmonary and lymph node granulomas across niches.

Figure S4

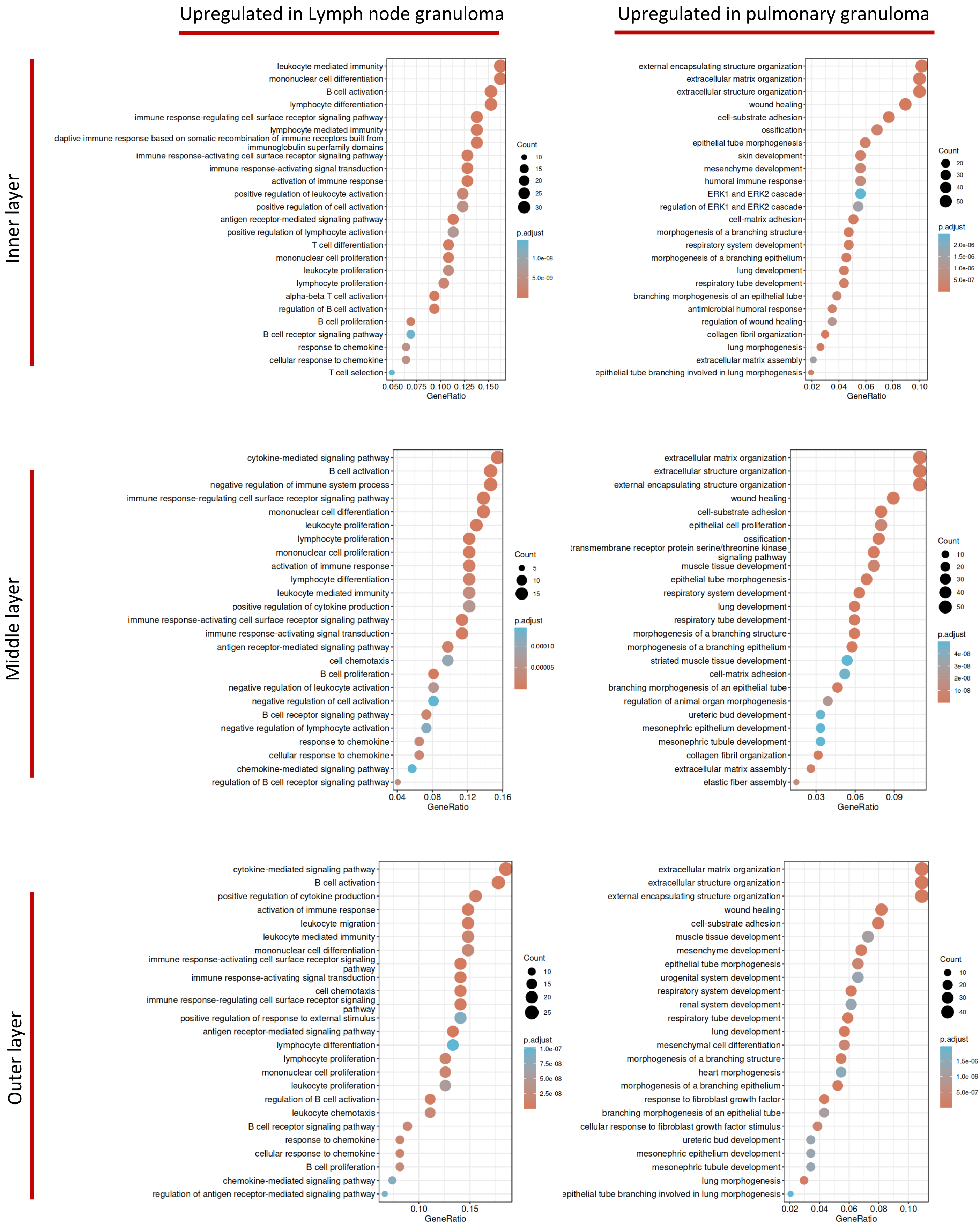


Supplementary Fig. 3
Biological process enrichment analysis of the DEGs in Fig. 4f in each niche.

Figure S5

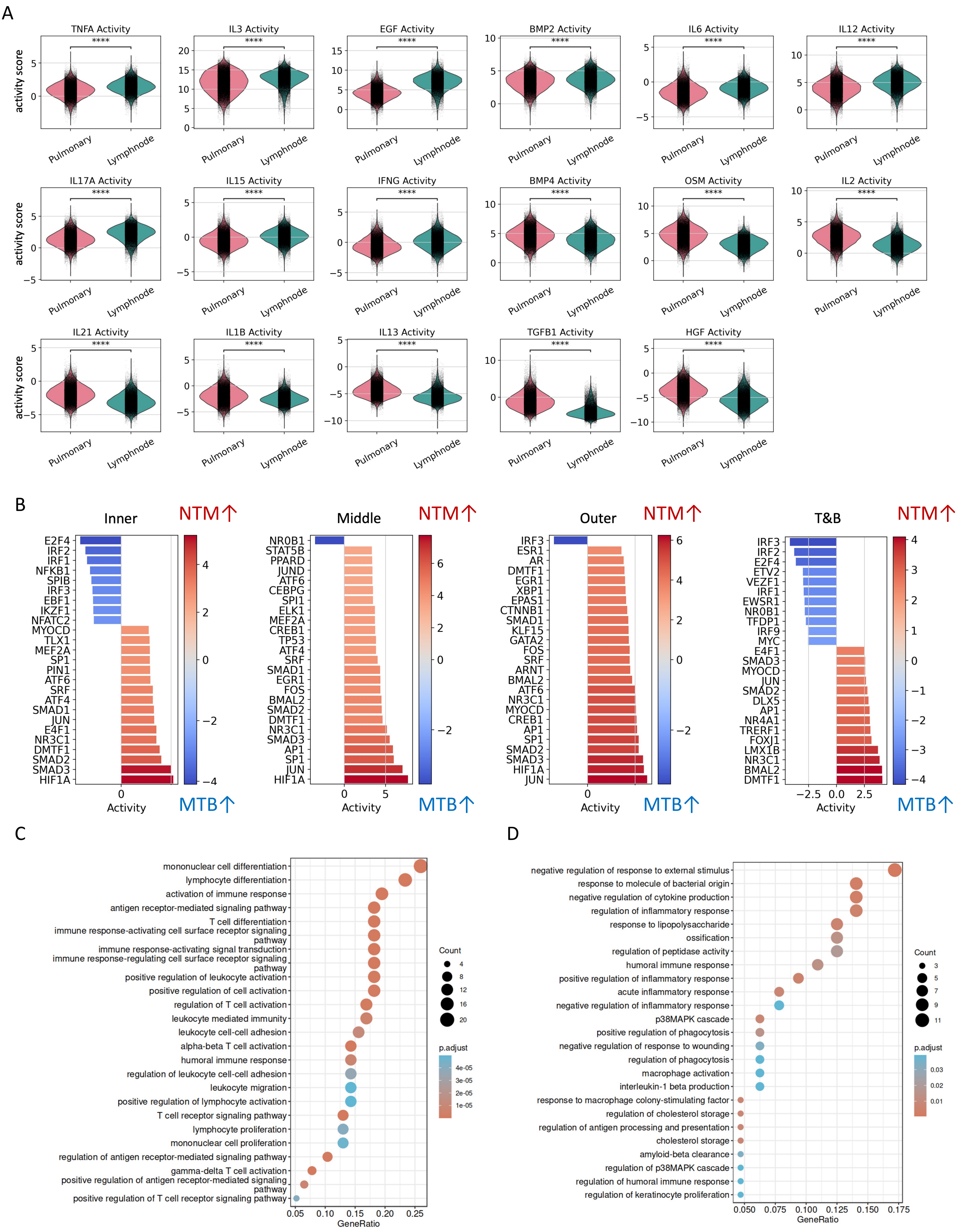


Supplementary Fig. 5
(a) Cytokine activity difference between pulmonary and lymph node granuloma in whole granuloma. (b) TF activity difference between NTM and MTB. (c) Biological process enrichment of the 81 upregulated genes in MTB granulomas, and (d) the 65 upregulated genes in NTM granulomas.

Figure S6

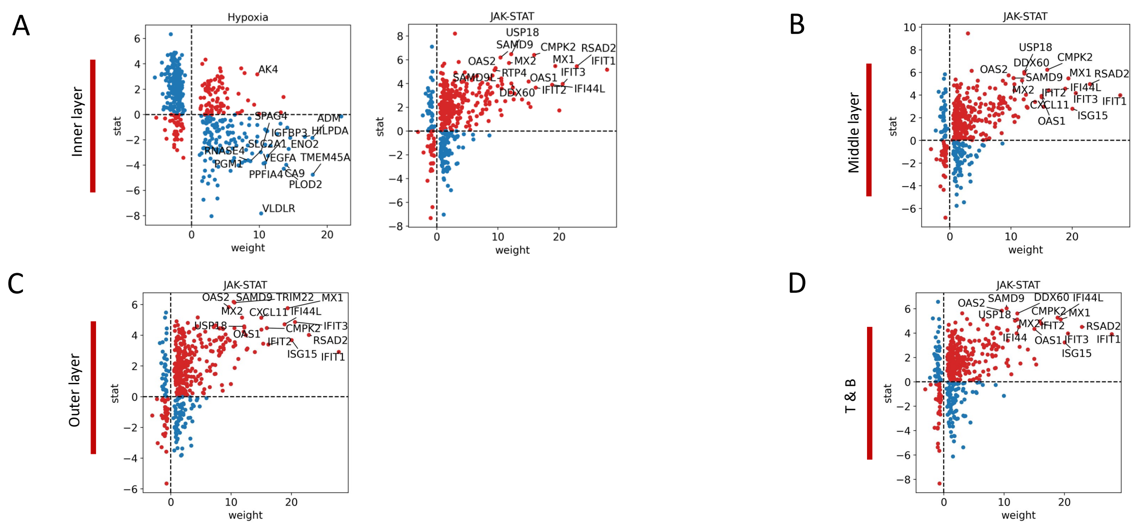


Supplementary Fig. 6. The DEGs make contribution to the differential pathway activity between Lymph node and Pulmonary granulomas.

(a) Expression level of maker genes for Hypoxia, and JAK-STAT pathway in the inner niche. (b) Expression level of maker genes for JAK-STAT pathway in the middle niche. (c) Expression level of maker genes for JAK-STAT pathway in the outer niche. (d) Expression level of maker genes for JAK-STAT pathway in the T&B niche. Red: genes make positive contribution to pathway activity in lymph node granulomas. Blue: genes make positive contribution to pathway activity in pulmonary granulomas.

Figure S7

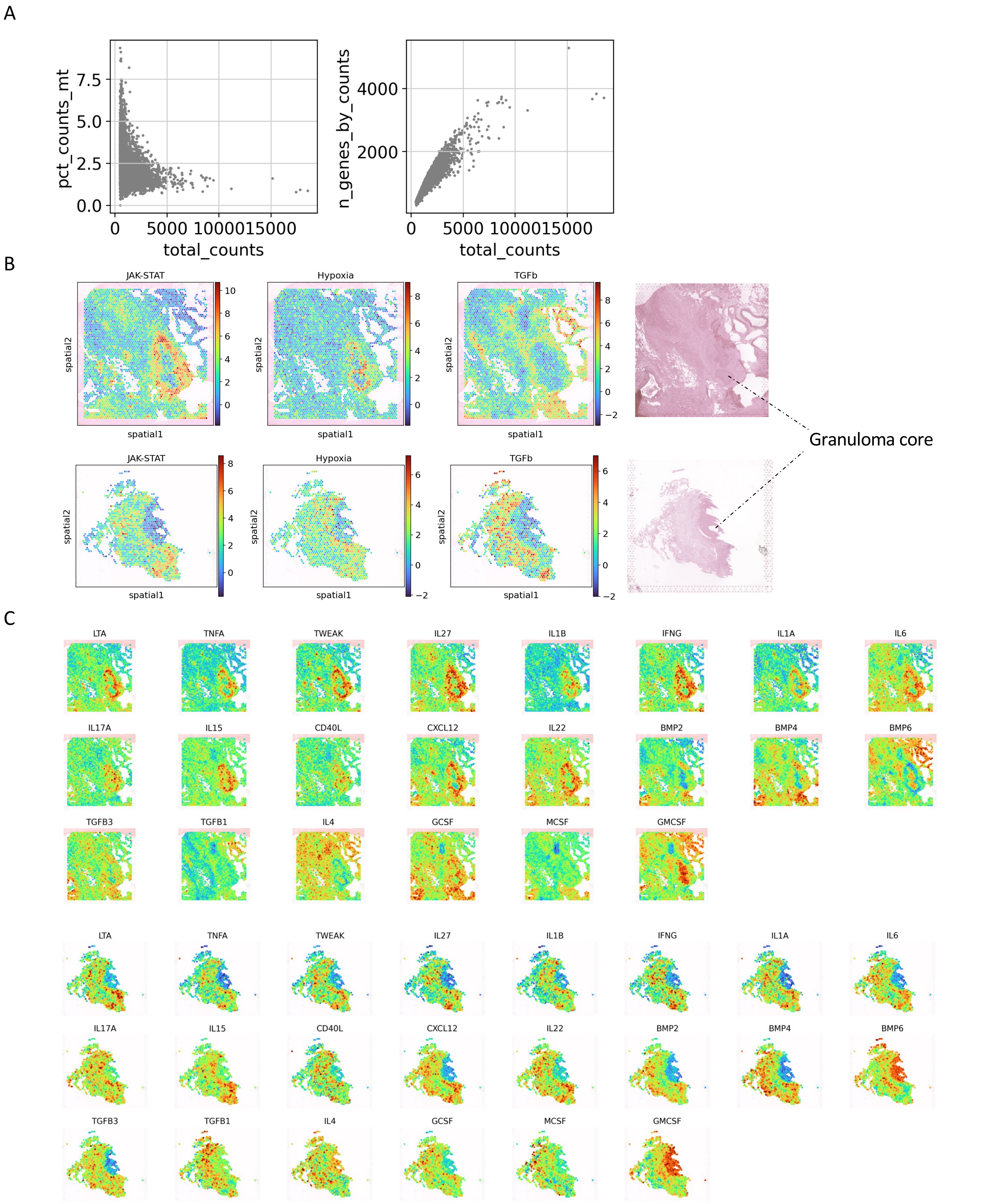


Supplementary Fig. 7

(a) Genes per spot and counts detected in the replication cohort. (b) Replication of the JAK-STAT, hypoxia, and TGFb pathway activity. (c) Cytokine activity replication.
